# Boltz-2: Towards Accurate and Efficient Binding Affinity Prediction

**DOI:** 10.1101/2025.06.14.659707

**Authors:** Saro Passaro, Gabriele Corso, Jeremy Wohlwend, Mateo Reveiz, Stephan Thaler, Vignesh Ram Somnath, Noah Getz, Tally Portnoi, Julien Roy, Hannes Stark, David Kwabi-Addo, Dominique Beaini, Tommi Jaakkola, Regina Barzilay

**Affiliations:** MIT CSAIL; MIT Jameel Clinic; Valence Labs; Recursion; ETH Zurich

## Abstract

Accurately modeling biomolecular interactions is a central challenge in modern biology. While recent advances, such as AlphaFold3 and Boltz-1, have substantially improved our ability to predict biomolecular complex structures, these models still fall short in predicting binding affinity, a critical property underlying molecular function and therapeutic efficacy. Here, we present Boltz-2, a new structural biology foundation model that exhibits strong performance for both structure and affinity prediction. Boltz-2 introduces controllability features including experimental method conditioning, distance constraints, and multi-chain template integration for structure prediction, and is, to our knowledge, the first AI model to approach the performance of free-energy perturbation (FEP) methods in estimating small molecule–protein binding affinity. Crucially, it achieves strong correlation with experimental readouts on many benchmarks, while being at least 1000*×* more computationally efficient than FEP. By coupling Boltz-2 with a generative model for small molecules, we demonstrate an effective workflow to find diverse, synthesizable, high-affinity binders, as estimated by absolute FEP simulations on the TYK2 target. To foster broad adoption and further innovation at the intersection of machine learning and biology, we are releasing Boltz-2 weights, inference, and training code ^1^ under a permissive open license, providing a robust and extensible foundation for both academic and industrial research.

## 1 Introduction

Complex biological processes are governed by interactions between biomolecules, including proteins, DNA, RNA, and small molecules. In this work, we introduce Boltz-2, a new foundation model for elucidating biomolecular interactions. Building on its predecessors AlphaFold3 [Abramson et al., 2024] and Boltz-1 [Wohlwend et al., 2025], Boltz-2 improves structural accuracy across modalities, extends predictions from static complexes to dynamic ensembles and sets a new standard in physical grounding. However, its key distinctive feature is its ability to predict binding affinity, which measures how tightly small molecules attach to proteins. This measure is critical for understanding whether a drug will act on its intended target and be potent enough to produce a therapeutic effect.

Despite its importance in drug design, in-silico affinity prediction remains an open challenge. To date, the most accurate techniques are atomistic simulations like free-energy perturbations (FEP). However, they are far too slow and expensive to be used at scale. Faster methods, such as docking, are not precise enough to give a reliable signal. In fact, no AI-based model has yet matched the accuracy of FEP methods or laboratory assays for binding affinity prediction.

Boltz-2 overcomes this long-standing performance/compute time trade-off. This advancement builds on two complementary developments: data curation and representation learning. Finding the right training signal for this task is a known barrier. While large amounts of binding data are publicly available, in their raw form they are not suitable for training due to experimental differences and noise. To this end, we standardized millions of biochemical assay measurements, tailoring data curation, sampling and supervision to extract the useful signal from the data.

In terms of representation learning, affinity prediction builds on the latent representation driving the cofolding process. This representation inherently encodes rich information about biomolecular interactions. Therefore, Boltz-2’s improvements in binding affinity prediction are driven by advances in structural modeling. These stem from: (1) extending training data beyond static structures to include experimental and molecular dynamics ensembles; (2) significantly expanding distillation datasets across diverse modalities; and (3) enhancing user control through conditioning on experimental methods, user-defined distance constraints, and multi-chain template integration.

The power of Boltz-2 to accurately predict affinity is evident in multiple discovery contexts:

- **Hit-to-lead and lead optimization** Boltz-2 significantly outperforms deep learning baselines on the FEP+ benchmark [Ross et al., 2023] and approaches the accuracy of FEP-based methods, while being over 1000 times faster (see Figure 1). On the CASP16 affinity track, retrospective evaluation shows that Boltz-2 outperforms all submitted competition entries out of the box.
- **Hit discovery** The model discriminates binders from decoys in high-throughput screens and achieves substantial enrichment gains on the MF-PCBA benchmark [Buterez et al., 2023], outperforming both docking and machine learning (ML) approaches.
- **De-novo Generation** Coupled with a generative model [Cretu et al., 2024], Boltz-2 enables discovery of new binders. In a prospective screening against the TYK2 target, this pipeline is able to generate diverse, synthetizable, high-affinity binders, as estimated by absolute binding free energy (ABFE) simulations [Wu et al., 2025].

**Figure 1:**
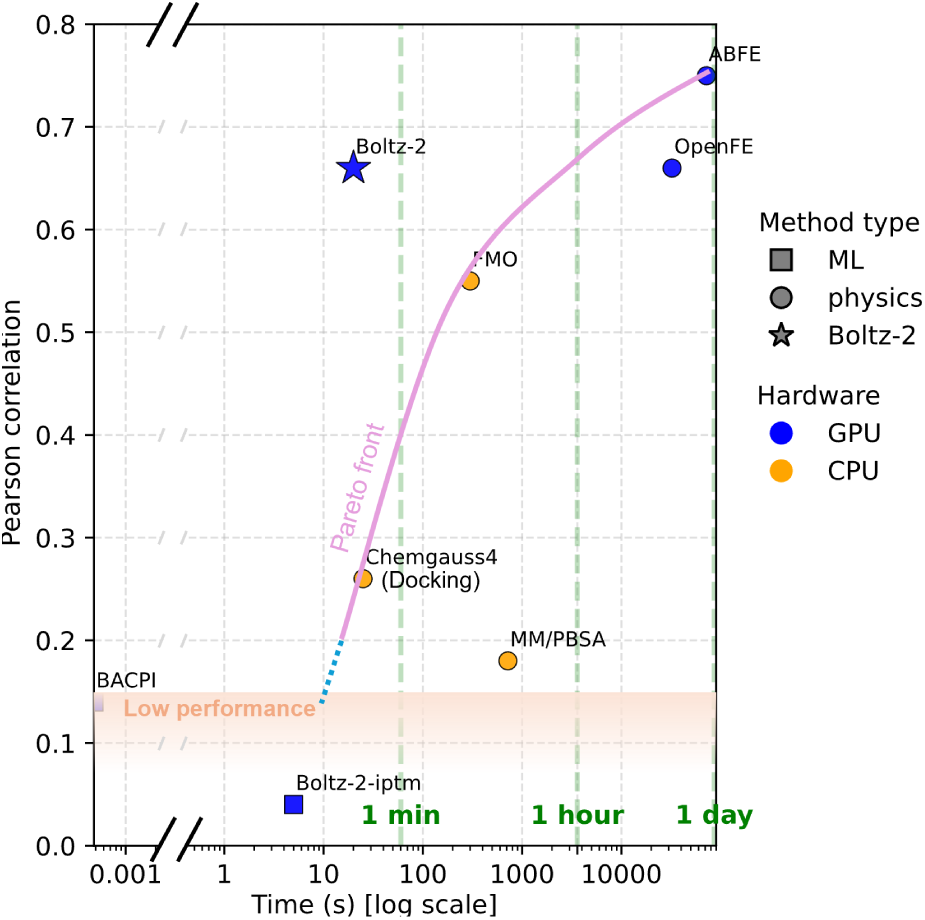
Boltz-2 presents a strong accuracy / speed trade-off for affinity prediction. Plot based on a 4-target subset (CDK2, TYK2, JNK1, P38) of the protein-ligand-benchmark [Hahn et al., 2022] for which baseline data are available for all methods. Full results in Figure 6.

Compared to Boltz-1, Boltz-2 improves crystallographic structure prediction across modalities, with notable gains on challenging targets such as antibody–antigen complexes. When benchmarked against molecular dynamics simulations, Boltz-2 matches the performance of recent specialized models, such as AlphaFlow [Jing et al., 2024] and BioEmu [Lewis et al., 2025], in predicting key dynamic properties like Root Mean Square Fluctuation (RMSF).

Alongside this manuscript, we are releasing Boltz-2’s model weights, inference pipeline and training code under a permissive open license. By making Boltz-2 freely available, we aim to accelerate progress across both academic and industrial efforts on tackling complex diseases and designing novel biomolecules. We also hope Boltz-2 will serve as a robust and extensible foundation for the growing machine learning community working at the interface of computation and biology, catalyzing further innovation in structure prediction, molecular design, and beyond.

## 2 Data

Aggregating and curating data are two of the most important steps in training strong foundational models. In this section, we summarize the training datasets and the key decisions made during data collection and preprocessing. Additional details are provided in Appendix A.

### Structural Data

For the structure model, we increased the diversity of biomolecules and data sources compared to Boltz-1. Unlike Boltz-1, which trained on a single structure per system, we supervise Boltz-2 using ensembles coming from both experimental techniques, such as NMR, as well as computational ones, such as molecular dynamics. The experimental data used for training comprises structures in the Protein Data Bank (PDB) [Berman et al., 2000] released before 2023-06-01. For molecular dynamics, we collected poses from the trajectories released as part of three large-scale open efforts: MISATO [Siebenmorgen et al., 2024], ATLAS [Vander Meersche et al., 2024], and md-CATH [Mirarchi et al., 2024]. Our goal is to expose Boltz-2 not only to single equilibrium points from crystal structures but also to local fluctuations and global structural ensembles.

To further improve the model’s understanding of local dynamics, we supervise the model’s single representation at the end of the trunk of the architecture to predict B-factors coming from both experimental methods as well as molecular dynamics trajectories.

In addition, we employ *distillation* to increase the size and diversity of the training data and its supervision signal. Distillation obtains additional training data by using high-confidence outputs of other models to augment the original training set. Specifically, we use AlphaFold2 high-confidence predictions on single-chain monomers [Varadi et al., 2022], like many previous models. Additionally, we employ high-confidence Boltz-1 prediction across a wide variety of complexes of single-chain RNA, protein-DNA, ligand-protein, MHC-peptide, and MHC-peptide-TCR interactions.

### Binding Affinity Data

Millions of binding affinity data points have been publicly released on central databases, such as PubChem [Kim et al., 2023] or ChEMBL [Zdrazil et al., 2024]; however, they have been notoriously difficult to combine into a single dataset for training due to variations in protocols and experimental noise [Landrum and Riniker, 2024].

Our data curation strategy focuses on: (1) retaining only the higher-quality assays, (2) mitigating overfitting to data biases by, for example, generating synthetic decoys, (3) ensuring structural quality by filtering targets with low confidence score, and (4) applying PAINS (pan-assay interference compounds) filters [Baell and Holloway, 2010] and discarding ligands with more than 50 heavy atoms.

Binding affinity predictions support two distinct tasks: hit discovery, where the goal is to identify likely binders across large chemical libraries, and hit-to-lead or lead optimization, where fine-grained affinity differences guide compound refinement. These use cases place different demands on the data: the former demands large-scale binary labeled data that distinguishes actives from inactives, while the latter requires precise, quantitative affinity measurements to resolve subtle activity differences. To support both settings, we curate a hybrid dataset comprising both binary and continuous labels. A summary of the resulting data is shown in Table 1.

**Table 1:** Summary statistics of the affinity training dataset used in our model. Each row corresponds to a different data source or curation strategy. The table reports the number of binders, decoys, unique protein clusters at 90% sequence identity (referred to as Targets in the table), and compounds. Supervision indicates whether the data is used to supervise the binary and/or affinity value head. Values in parentheses show the corresponding statistics prior to applying the structural quality filter, which excludes examples with iptm below 0.75.

| Source | Type | Supervision | # Binders | # Decoys | # Targets | # Compounds |
| --- | --- | --- | --- | --- | --- | --- |
| ChEMBL and BindingDB | optimization | values | 1.2M (1.45M) | 0 | 2k (2.5k) | 600k (700k) |
| PubChem small assays | hit-discovery | both | 10k (13k) | 50k (70k) | 250 (300) | 20k (25k) |
| PubChem HTS | hit-discovery | binary | 200k (400k) | 1.8M (3.5M) | 300 (500) | 400k (450k) |
| CeMM Fragments | hit-discovery | binary | 25k (45k) | 115k (200k) | 1.3k (2.5k) | 400 (400) |
| MIDAS Metabolites | hit-discovery | binary | 2k (3.5k) | 20k (35k) | 60 (100) | 400 (400) |
| ChEMBL and BindingDB | synthetic decoys | binary | 0 | 1.2M (1.45M) | 2k (2.5k) | 600k (700k) |

For the **binding affinity regression** values (e.g., *K_i_*, *K_d_*, *IC*_50_, *AC*_50_, *EC*_50_, *XC*_50_), we gather data from PubChem [Kim et al., 2023], ChEMBL [Zdrazil et al., 2024], and BindingDB [Liu et al., 2007]. We retain only assays that target a single protein and are categorized as either biochemical or functional, excluding any labeled as low-confidence or unreliable. All affinity values are standardized to log_10_ scale derived from values measured in *µ*M. Assays with insufficient data or a low affinity standard deviation are discarded to encourage learning of intra-assay differences in values rather than inter-assay.

For the **binary affinity classification** data, we gather data from PubChem HTS (high-throughput screening) assays [Kim et al., 2023], a fragment screening dataset from CeMM [Offensperger et al., 2024], and MIDAS, a protein–metabolite interactome dataset from the University of Utah [Hicks et al., 2023]. For PubChem HTS, we retain only assays that include at least 100 compounds and exhibit a hit rate below 10%, helping to filter out noisy screens. To reduce false-positive labels introduced by HTS noise, we check for the presence of an associated quantitative affinity measurement (e.g., *K_i_*, *K_d_*, or *XC*_50_) in independent assays. Lastly, we augment the binary classification dataset by generating **synthetic decoys** created by shuffling binders identified in hit-to-lead screens across different targets, while mitigating low false negative rates by ensuring that each decoy has a Tanimoto similarity below 0.3 to all known binders associated with similar proteins. This expands the pool of negative examples, improves coverage of the chemical space surrounding each protein target, and helps mitigate spurious correlations present in HTS assays.

## 3 Architecture

As shown in Figure 2, Boltz-2’s architecture comprises four main components: the trunk, the denoising module with additional steering components, the confidence module, and the affinity module. Below, we highlight the major differences compared to the Boltz-1 and Boltz-1x architectures, mostly related to the controllability components and the affinity module. Appendix B provides a detailed description of each component.

**Figure 2:**
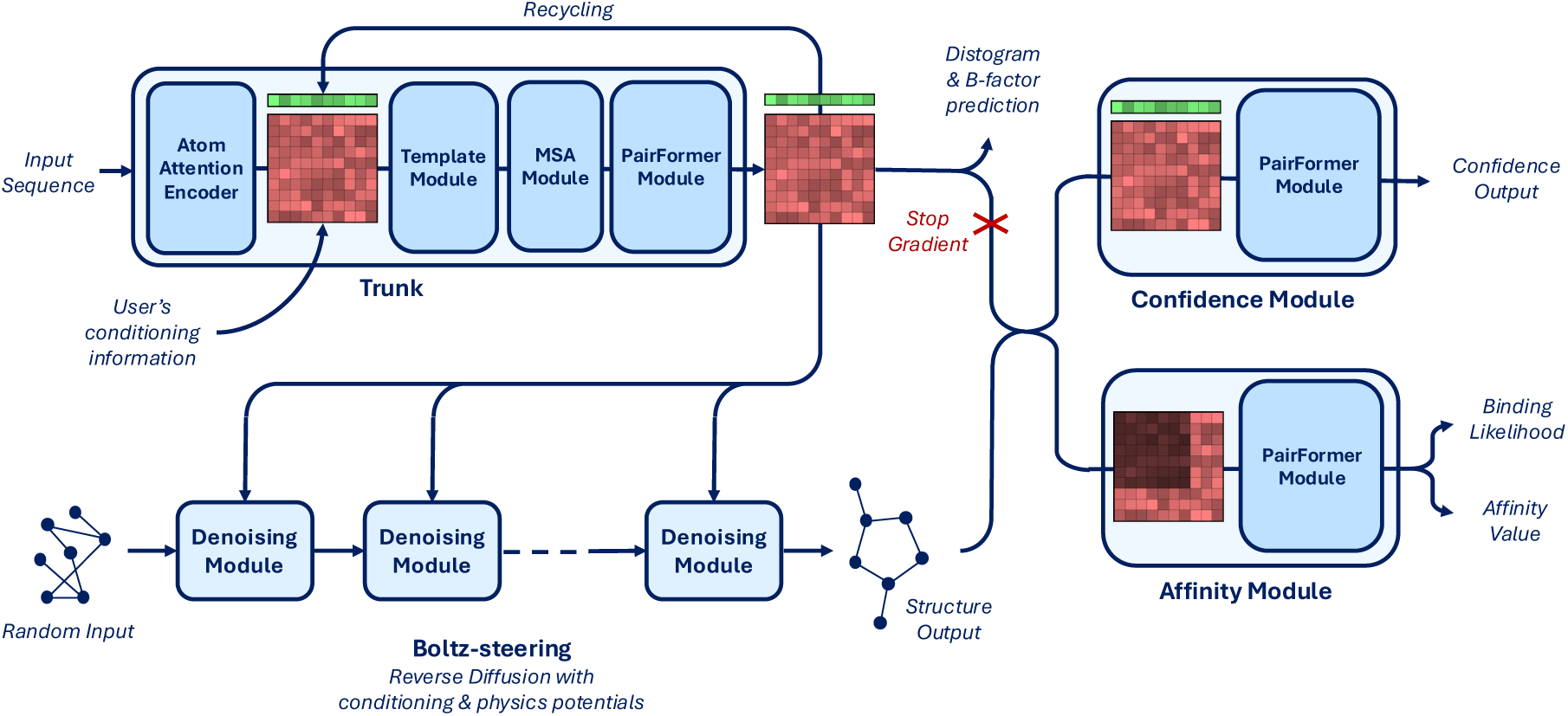
Boltz-2 model architecture diagram.

### Trunk optimization

The trunk is the most resource-intensive component of the model, largely due to the pairwise stack and triangular operations. We significantly improve the training and inference runtime as well as its memory consumption by using mixed-precision (bfloat16) and the trifast kernel for triangle attention. This also allows us to scale the crop size during training to 768 tokens, as done by AlphaFold3.

### Physical quality

Co-folding models such as AlphaFold3, Chai-1, and Boltz-1 often produce structures with physical inaccuracies such as steric clashes and incorrect stereochemistry [Abramson et al., 2024, Buttenschoen et al., 2024]. To address this, we recently introduced Boltz-steering (as part of the Boltz-1x release) — an inference-time method that applies physics-based potentials, which improves physical plausibility without sacrificing accuracy. We also integrate this approach within Boltz-2 to obtain Boltz-2x.

### Controllability

A frequent request from Boltz-1 users was a desire for more precise control of the model’s predictions, allowing them to test hypotheses or incorporate prior knowledge into the model without costly retraining or fine-tuning. To enable better controllability of the poses, we integrate three new components in Boltz-2: *method conditioning*, *template conditioning and steering*, and *contact and pocket conditioning*. Method conditioning allows for specification of the type of structure prediction method (e.g., X-ray crystallography, NMR, or molecular dynamics) that the predictions should align with and can capture their many nuances (see Section 5.2). Template conditioning integrates structures of similar complexes, helping the model without retraining [Jumper et al., 2021]. Unlike previous approaches, we allow users to either enforce strict observance of the templates via steering or just use the soft-conditioning like previous methods. As a departure from previous work, our templating approach also natively supports the use of multimeric templates. Finally, contact and pocket conditioning allow for specification of particular distance constraints, whether they come from experimental techniques or human intuition.

### Affinity module

The affinity module consists of a PairFormer and two heads: one predicting binding likelihood, the other regressing continuous affinity values. During training, we supervise the affinity value head using a mixture of related, but non-identical biochemical quantities (including *K_i_*, *K_d_*, and *IC*_50_) all converted to the logarithmic scale using *µ*M as standardized unit. While some of these measures are related through the Cheng–Prusoff equation, they arise from different experimental contexts. As such, the predicted value should be viewed as a general measure of binding strength that supports ranking and can be approximately interpreted as an *IC*_50_-like value. The module operates on Boltz-2’s structural predictions, leveraging the pair representation and the predicted coordinates refined by a PairFormer model focused exclusively on the protein–ligand and intra-ligand interactions. These interactions are then aggregated to produce both a binding likelihood and an affinity value.

## 4 Training

The training of the model can be divided into three phases: structure training, confidence training, and affinity training. We further discuss how we use Boltz-2 to train a generative model for efficient exploration of the synthesizable chemical space. Full details on these components can be found in Appendix C.

### Structure and Confidence training

The structure and confidence training largely follows Boltz-1, with a few exceptions. (1) Computational optimizations allowed us to train the model for more iterations and larger crops. (2) Ensembles from experimental methods and molecular dynamics were supervised with an aggregated distogram to reduce variance. (3) The trunk’s final representation was also supervised to predict the B-factor of each token.

### Affinity training

Affinity training is performed after structure and confidence training, with gradients detached from the trunk. The pipeline incorporates several key components designed to improve generalization and scalability: pre-computation and cropping of binding pockets to focus on the most relevant interactions, pre-processing of trunk representations and a custom sampling strategy that balances binders and decoys while prioritizing informative, high-contrast assays. Batches are constructed to focus on local chemical variation. Supervision is applied jointly across binary and continuous affinity tasks using robust loss functions designed to mitigate the effects of experimental noise and assay heterogeneity. Continuous values are supervised using a Huber loss applied to both **absolute** affinity values and, with stronger weight, to the **pairwise** intra-assay differences. We observed best performance when training a single affinity value head on all available affinity measurements (eg, *K_i_*, *K_d_*, *IC*_50_, *AC*_50_, *EC*_50_, and *XC*_50_). Although these metrics reflect different underlying biochemical quantities, *K_i_*s and *IC*_50_s are related through the Cheng–Prusoff equation, and when comparing affinity values within the same assay, the **pairwise** differences loss effectively cancel out the correction term, and assays can be combined Ross et al. [2023]. Binary classification is supervised using a focal loss [Lin et al., 2017] to address class imbalance and reduce overfitting. The final training objective is a weighted combination of the classification and the regression losses, designed to balance the different tasks.

### Training a molecular generator with Boltz-2

As part of our evaluation, Boltz-2 is used to train a molecular generator to produce small molecules with high binding scores. Our generative agent (SynFlowNet [Cretu et al., 2024]) employs a GFlowNet [Bengio et al., 2021] loss function, enabling it to sample from arbitrary and multi-modal score distributions. Within this framework, the molecular generator undergoes off-policy training: batches of candidate molecules are asynchronously submitted to Boltz-2 workers for scoring, and the results are then incorporated into a replay buffer for the generative agent. The binding score (reward) for the agent is a strictly positive metric derived from a combination of both the binding likelihood and affinity values predicted by Boltz-2. The training procedure also incorporates basic drug-likeness properties through medicinal chemistry filters.

## 5 Evaluation

In this section, we evaluate Boltz-2 in various settings, including crystal structure prediction, local protein dynamics, binding likelihood and affinity predictions, and virtual screening. For the affinity measurements, all cross-assay averages are weighted by the number of compounds in the assay and error bars are computed as the bootstrapping standard deviation.

### 5.1 Boltz-2 improves over Boltz-1 on structure prediction

#### PDB evaluation set

We evaluated the performance of Boltz-2 (and its version with enabled physicality steering potentials, Boltz-2x), comparing it against Boltz-1, Chai-1 [Chai et al., 2024], ProteinX [Chen et al., 2025], and AlphaFold3 and across a wide variety of complexes submitted to the Protein Data Bank in 2024 and 2025 that were significantly different from any structure that any of the models had seen in their training set. The results, presented in Figure 3, show that, across modalities, Boltz-2 matches or moderately improves over the performance of Boltz-1. Among the modalities where the improvements are strongest are RNA chains and DNA-protein complexes. These are the two modalities where we most significantly augmented the available data in the PDB with large distillation sets, suggesting that the distillation strategy could be important to improve these models beyond what available experimental data allows. Compared also to other methods, Boltz-2 performs competitively, edging the other commercially available models Chai-1 and ProteinX, but lagging a bit behind AlphaFold3. As expected Boltz-1x and Boltz-2x, thanks to Boltz-steering obtain significantly better physicality metrics both for small-molecule conformations and for steric clashes at interfaces.

**Figure 3:**
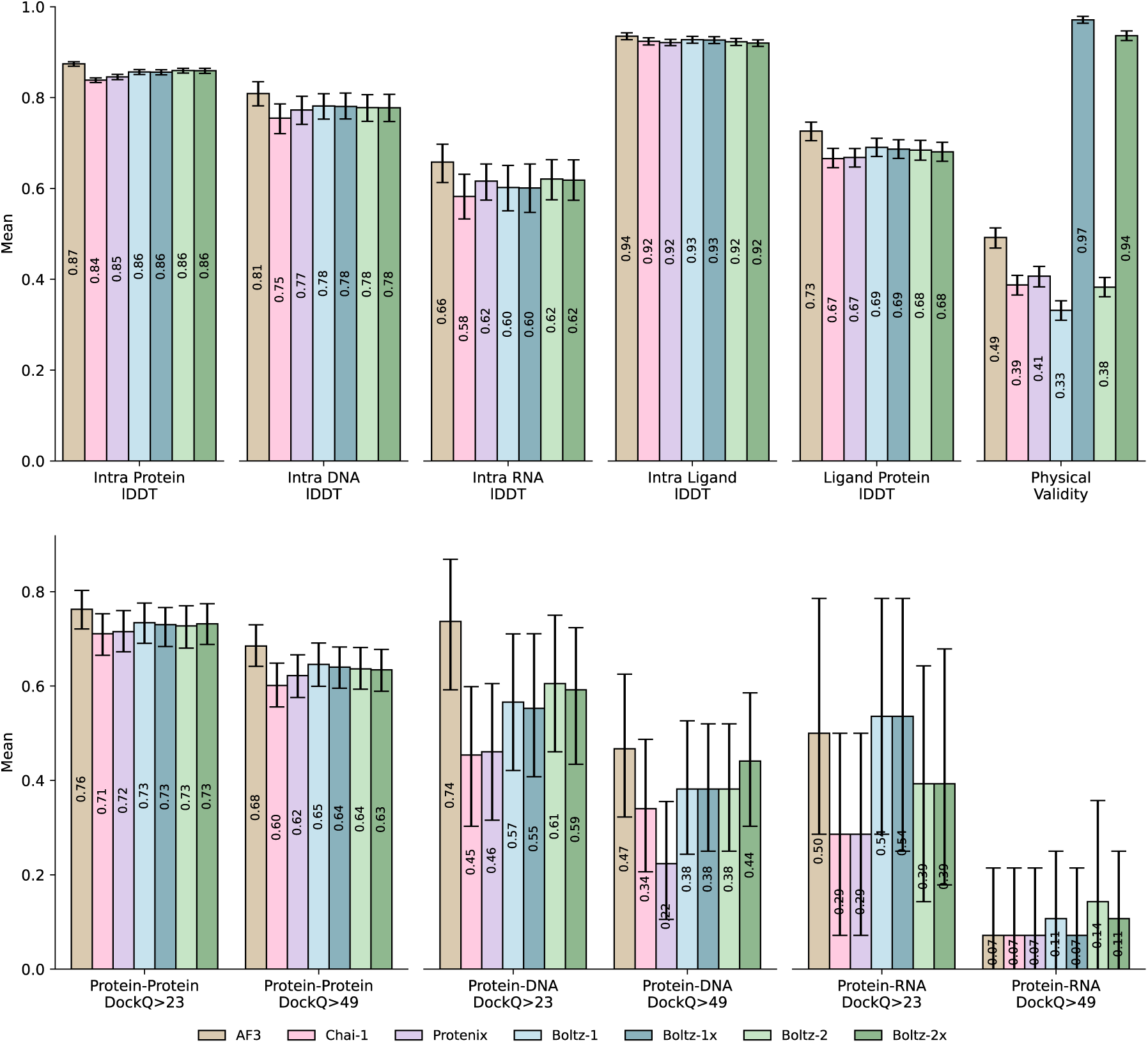
Evaluation of the performance of Boltz-2 against existing co-folding models on a diverse set of unseen complexes. Error bars indicate 95% confidence intervals.

#### Antibody benchmark

One modality where researchers have highlighted a performance gap between AlphaFold3 and the commercially-available models is antibody-antigen structure prediction, especially when looking at the generalization to unseen antigens. This observation is also reflected in the results from our antibody benchmark shown in Figure 4. However, we also observe a moderate improvement of Boltz-2 over Boltz-1, narrowing the gap between the proposed open models and proprietary ones, such as AlphaFold3.

**Figure 4:**
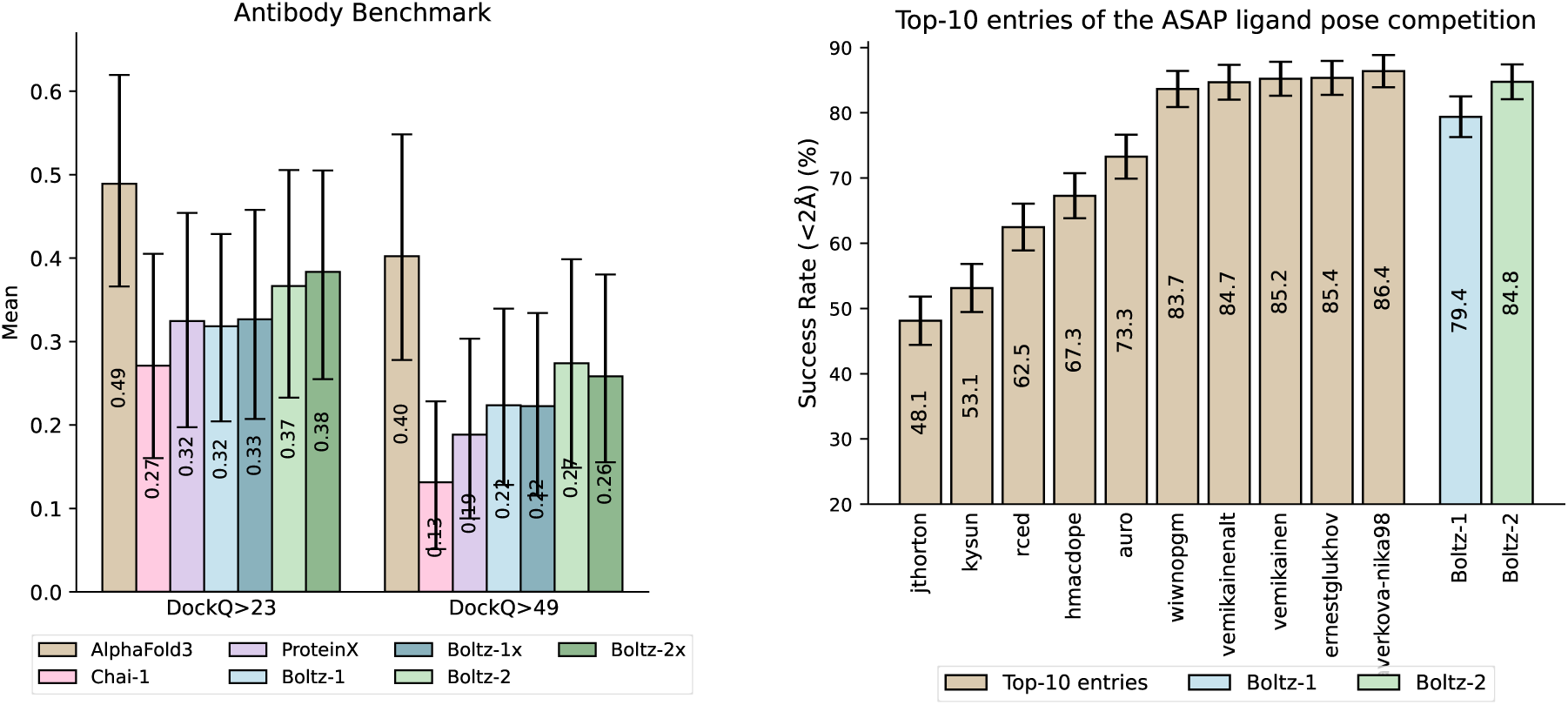
*Left:* Performance of different co-folding methods on a challenging antibody benchmark. Boltz-2 shows an improvement over Boltz-1 while still lagging behind AlphaFold3. *Right:* Retrospective results for the Polaris-ASAP competition, with Boltz-2 matching the performance of the top 5 contenders without any fine-tuning or physics relaxation. Error bars indicate 95% confidence intervals.

#### Polaris-ASAP challenge (SARS-CoV-2 and MERS-CoV)

We further evaluated the model on the recent Polaris-ASAP Discovery competition on ligand pose estimation. This was composed of ligands bound to either the SARS-CoV-2 and MERS-CoV main proteases that ASAP Discovery generated as part of their antiviral drug discovery campaigns. On top of the PDB, 770 additional structures of similar ligands bound to these proteins were given as a training set. This challenge saw a clear success of co-folding models over more traditional physics-based and ML tools, with all the top-6 entries being composed of fine-tuned Boltz-1 or AlphaFold3 models (some with additional physics-based relaxation). Boltz-2 shows a clear improvement over Boltz-1 and the top performers in the challenge, without any finetuning or physics-based relaxation (Figure 4 right).

### 5.2 Boltz-2 can better capture local protein dynamics

In order to validate the impact of MD method conditioning and evaluate the model’s ability to capture local dynamics of protein structures, we evaluated Boltz-2 on the held-out clusters of the mdCATH and ATLAS datasets. The results, presented in Figure 5 and Appendix E.1, show that (1) MD conditioning has a clear effect on the predicted ensembles, leading to more diverse structures that better capture the conformational diversity of the simulations, (2) Boltz-2 with MD conditioning is competitive on various metrics with specialized models such as BioEmu [Lewis et al., 2025] and AlphaFlow [Jing et al., 2024]. When looking at RMSF, a standard measure of local dynamics, Boltz-2 MD ensembles generally obtain stronger correlations with the ground truth simulation and lower errors than Boltz-1, BioEmu and AlphaFlow. In addition to training on MD ensembles, Boltz-2’s performance may also benefit from supervision on both experimental and computational B-factor estimates, which are specifically designed to capture local structural dynamics. Looking at recall lDDT, Boltz-2 modestly outperforms Boltz-1 while improving over AlphaFlow and BioEmu. Conditioning on MD allows Boltz-2 to increase the diversity of samples while retaining its precision. This diversity increase is, however, outperformed by BioEmu and AlphaFlow, which more closely align with the reference diversity from the simulation.

**Figure 5:**
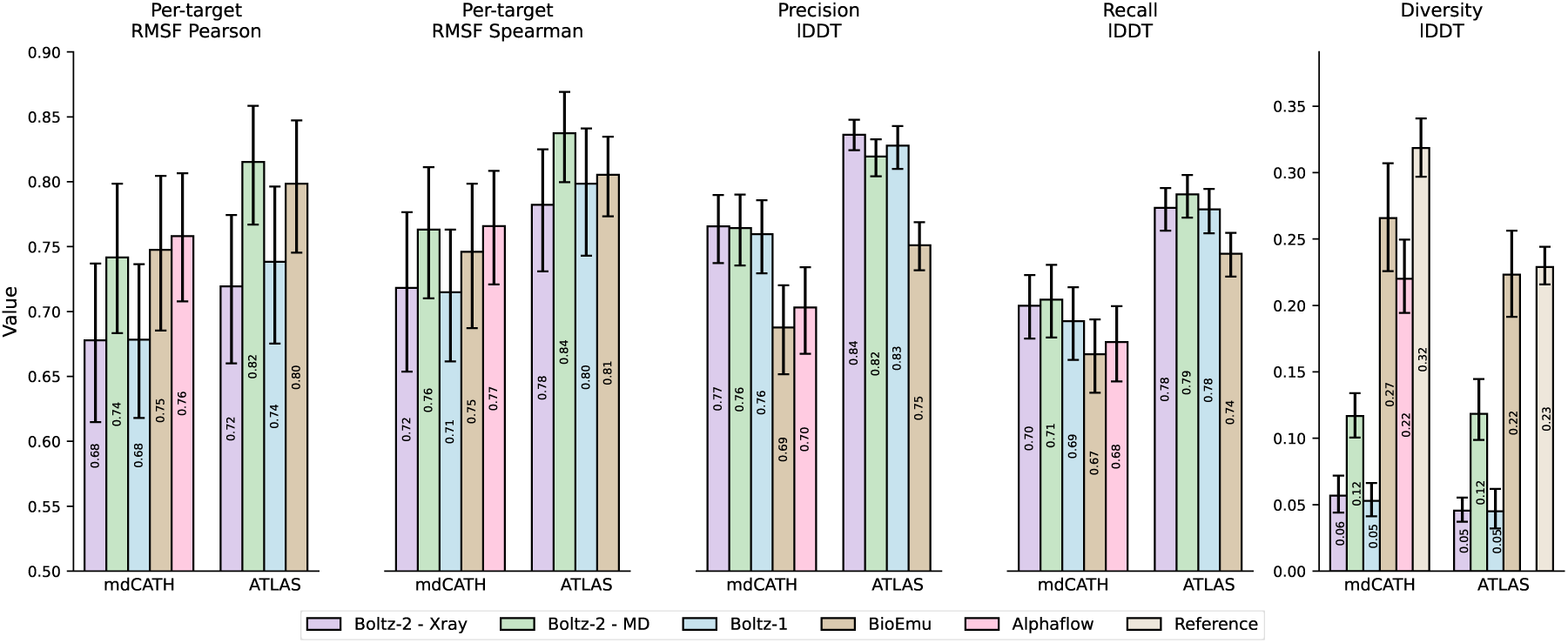
Per-target RMSF correlation metrics and lDDT metrics on the held-out clusters from the mdCATH and ATLAS molecular dynamics datasets.

### 5.3 Boltz-2 approaches FEP accuracy on public benchmarks

Accurately ranking analogues within a chemical series is a critical challenge in hit-to-lead and lead optimization. Distinguishing subtle differences in binding affinity among closely related analogues is essential for guiding molecular refinement and progressing candidates through the pipeline. Traditional free energy simulation methods can often offer the required precision, but are too computationally expensive for more widespread use. Boltz-2 addresses this problem as it allows accurate affinity predictions at a fraction of the computational cost, enabling rapid prioritization in structure-guided optimization workflows.

To evaluate Boltz-2’s affinity prediction ability, we benchmarked it across a suite of hit-to-lead and lead-optimization datasets. Summary results are presented in Figure 6, while expanded tables and scatter plots are available in Appendices E.2.1 and E.2.2.

**Figure 6:**
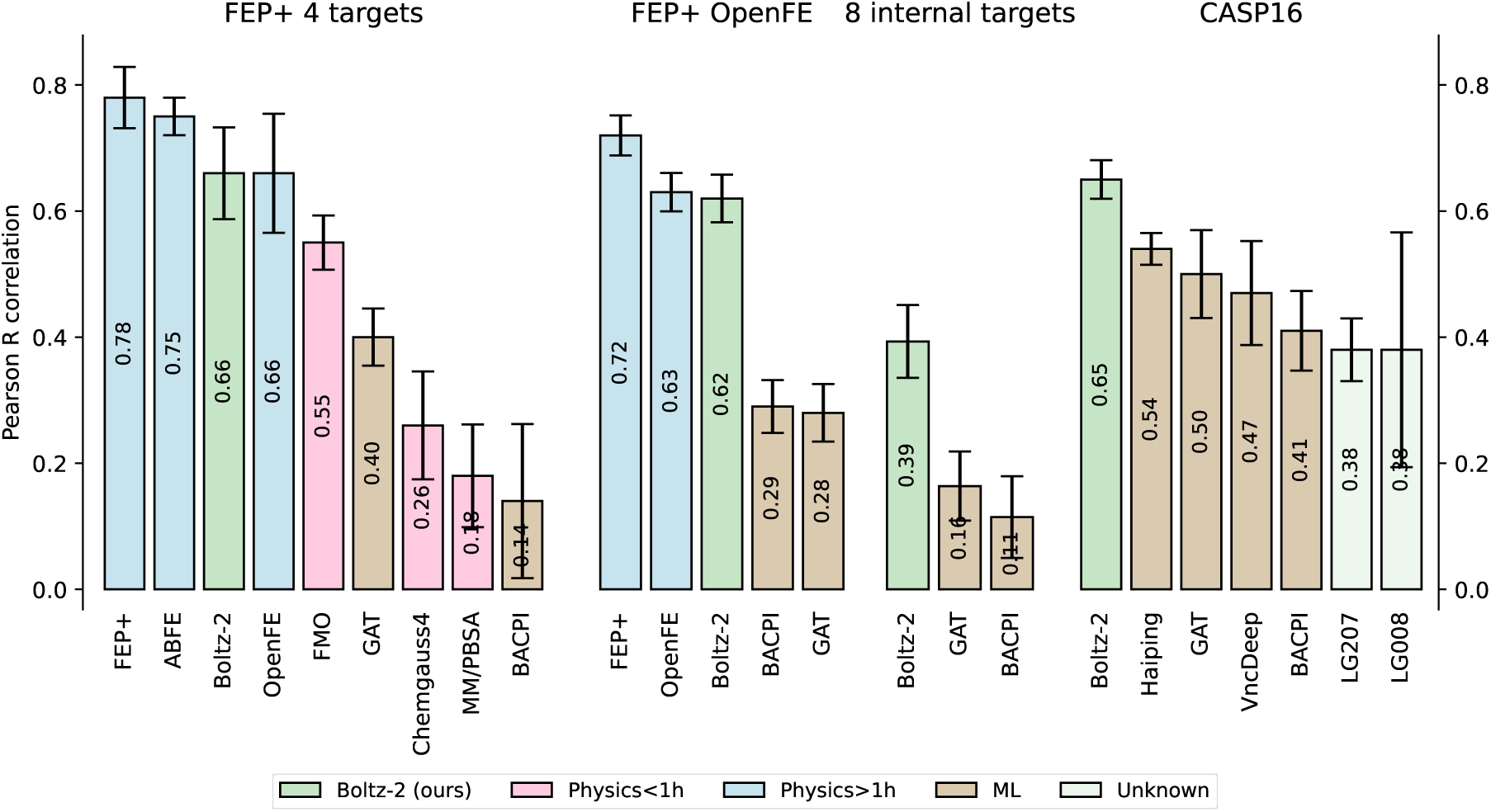
Pearson correlation averaged over each assay on our four affinity value test sets. Error bars represent bootstrap estimates of the standard error.

We evaluate the model on two subsets of the FEP+ benchmark [Ross et al., 2023]: the OpenFE dataset, consisting of 876 high-quality hit-to-lead measurements [Gowers et al., 2023], and a focused 4-target subset [Hahn et al., 2022], where more physics-based baselines are available, including absolute FEP (ABFE) [Wu et al., 2025] and Fragment Molecular Orbital (FMO) [Nishimoto and Fedorov, 2016, Guareschi et al., 2023], a semi-empirical quantum mechanics-based scoring function. The training sets are filtered to exclude proteins with ≥ 90% sequence identity to any protein in the FEP+ benchmark, ensuring that we benchmark on unseen proteins. Additionally, we assess the impact of compound similarity in Figure D.2.1. On the 4-target FEP subset, Boltz-2 achieves an average Pearson correlation of 0.66, outperforming all available inexpensive physical methods and ML baselines. Remarkably, Boltz-2 approaches state-of-the-art free energy simulations, while running more than 1,000× faster, providing a strong speed-accuracy tradeoff (Figure 1). Even on the full OpenFE benchmark set, Boltz-2 approaches the performance of OpenFE, a widely adopted open-source relative FEP method.

Additionally, we include the CASP16 affinity challenge [Gilson et al., 2025], a rigorous blind benchmark featuring 140 protein–ligand pairs across two targets. Here, while participants were given several weeks and used a range of ad-hoc machine learning and physics-based tools, we ran Boltz-2 out-of-the-box with no fine-tuning or input curation. Yet, Boltz-2 outperforms all top-ranking participants by a clear margin.

We also evaluated the model on eight blinded internal assays from Recursion that reflect complex real-world medicinal chemistry projects. Here, the model still outperforms by a large margin the other ML baselines and achieves a Pearson correlation of *>* 0.55 on 3 out of 8 assays, but has limited performance on the other 5. Such variation is also typical of FEP methods, which are known to perform weakly on some protein classes, such as GPCRs, without custom input preparation [Deflorian et al., 2020]. We include these results as a reminder that strong performance on public benchmarks does not always immediately translate to all complexities of real-world drug discovery without further work to understand the relative strengths and weaknesses of a given approach.

### 5.4 Boltz-2 enables accurate large-scale virtual screening

Accurate virtual screening remains one of the most impactful challenges in early-stage drug discovery. The ideal method must scale across vast chemical libraries while reliably identifying active compounds against diverse protein targets. Boltz-2 offers a promising solution to this problem, combining speed and precision in a unified affinity prediction framework.

To assess its utility in realistic screening settings, we first evaluated Boltz-2 on retrospective benchmarks derived from the MF-PCBA dataset [Buterez et al., 2023], which includes high-quality biochemical assays spanning diverse protein families. Performance was assessed using metrics tailored to hit discovery—average precision (AP), enrichment factor at top-ranked percentiles, and AUROC. Results highlight Boltz-2’s ability to retrieve actives from large, imbalanced datasets (Figure 7). On this benchmark, Boltz-2 substantially outperforms prior machine learning approaches, the widely used ipTM and docking, nearly doubling the average precision and achieving an enrichment factor of 18.4 at a 0.5% threshold (Table 13).

**Figure 7:**
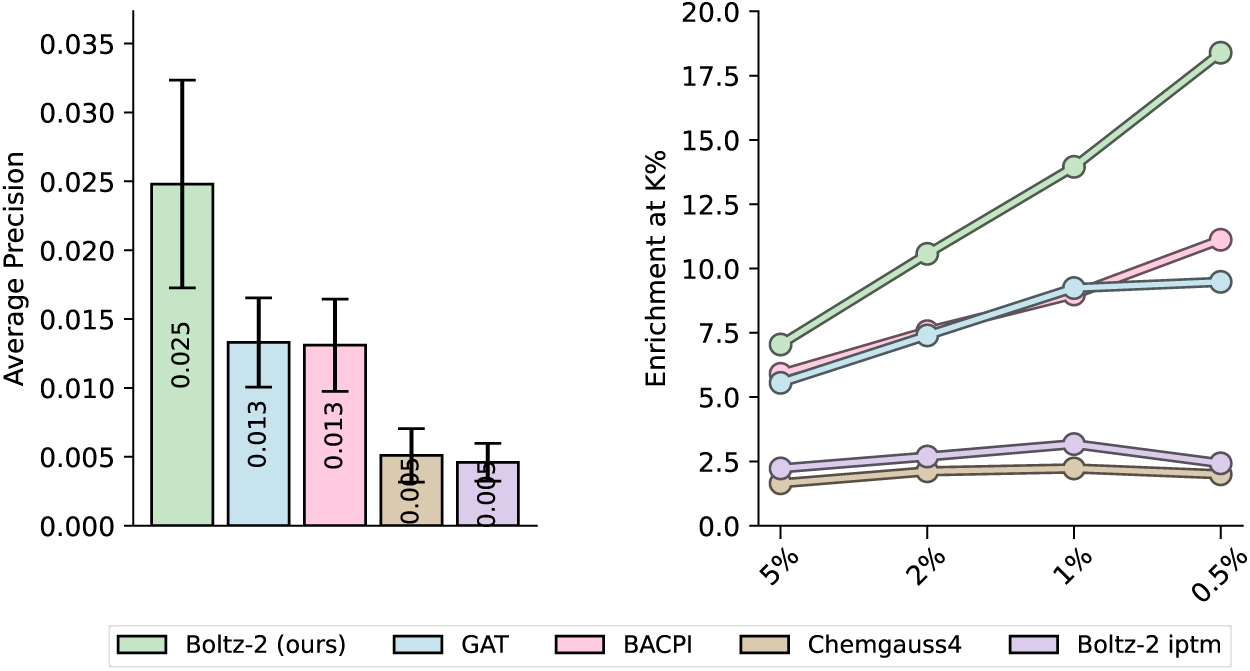
*Left:* Average precision averaged over the assays in the MF-PCBA test set. Error bars represent bootstrap estimates of the standard error. *Right:* Enrichment factors, computed at top-K thresholds with K = 0.5%, 1%, 2%, and 5%.

To evaluate Boltz-2 in prospective settings, we performed a virtual screen against the kinase target TYK2, a protein well-characterized in both ML and physics-based modeling benchmarks. We selected TYK2 for two main reasons: First, TYK2 is in the test set of the Boltz-2 affinity model, avoiding data leakage from known binders. Second, in the absence of experimental data, we validate the compounds selected by Boltz-2 with a single repeat of Boltz-ABFE^2^ [Wu et al., 2025], our recently developed absolute FEP pipeline to estimate ABFE values without experimental crystal structures, and Boltz-ABFE performs very well on this target. Indeed, based on the protein-ligand benchmark [Hahn et al., 2022], Boltz-ABFE achieves a Pearson R = 0.95, centered MAE = 0.42 kcal/mol and a comparatively small offset with respect to the experiment of 0.92 kcal/mol, supporting our confidence of this procedure as a validation step for TYK2-targeting virtual screens.

In these screens, we use a combination of the Boltz-2 predicted binding likelihood and affinity as a screen score for small molecules. We started by screening two commercially available compound libraries from Enamine—Hit Locator Library (HLL, 460,160 compounds) and Kinase Library (64,960 compounds). Boltz-2 successfully prioritized high-affinity ligands: Based on ABFE estimates, 8 of the top 10 compounds from HLL and all 10 compounds from the Kinase library are predicted to bind, while all 10 random compounds are predicted to be non-binders (Figure 8).

**Figure 8:**
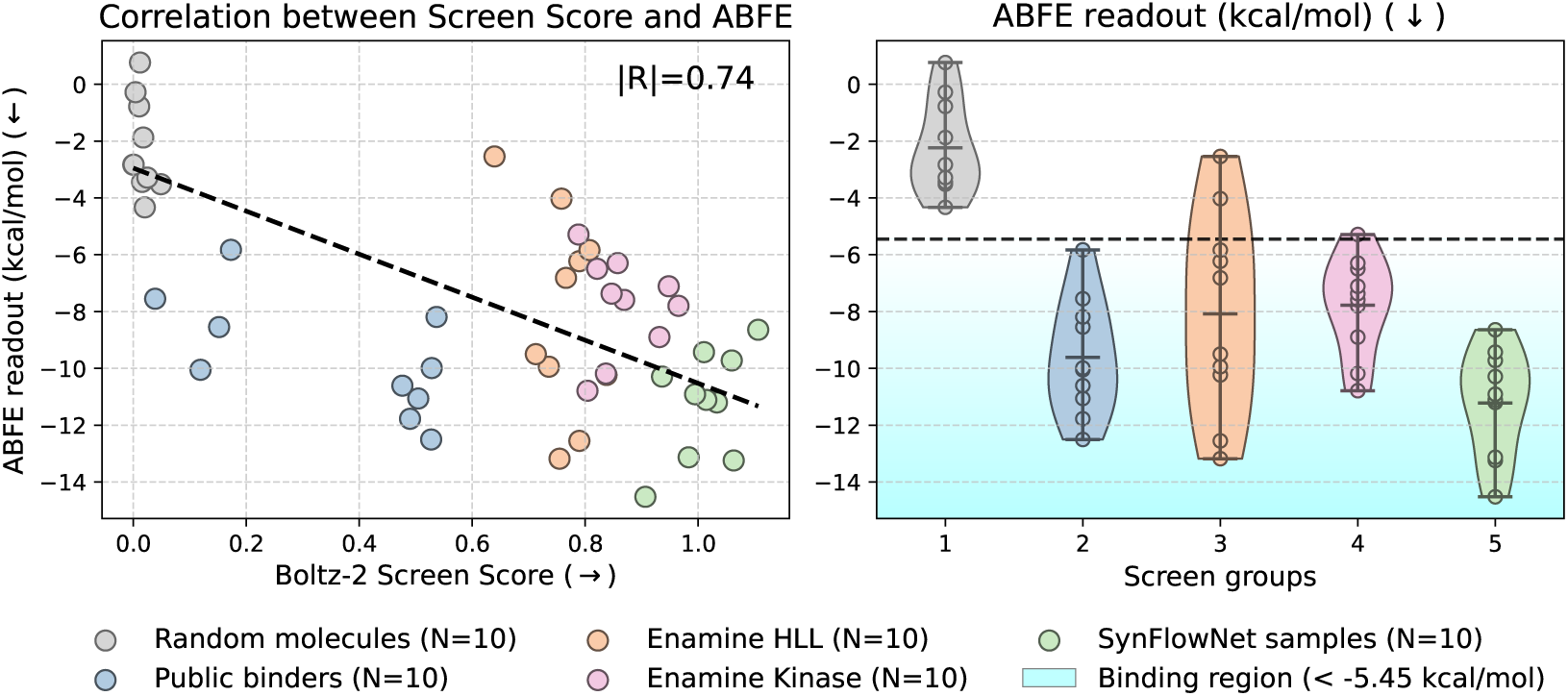
Virtual screening experiment performed on the TYK2 protein. *Left* : The Boltz-2 screen scores of the final set of compounds of each virtual screening stream correlate (|*R*| = 0.74) with the absolute binding free energy (ABFE) estimates Δ*G*. *Right* : Distribution of the ABFE-predicted Δ*G* for the compounds proposed by the different screening strategies.

**Figure 9:**
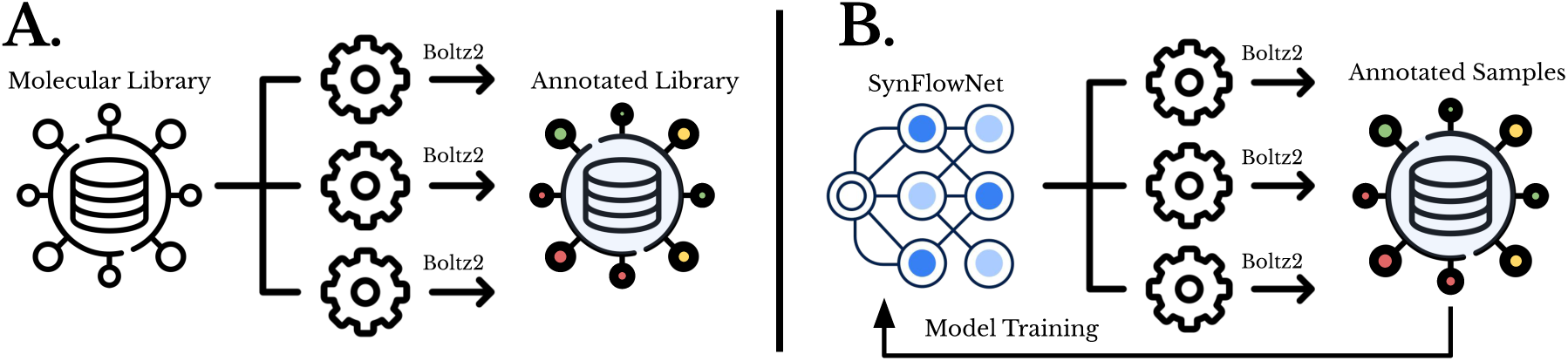
Depiction of two virtual screening strategies. A) Fixed-Library Virtual Screen: We start from a fixed molecular library and leverage high-performance computing infrastructure to accelerate the screening time by employing multiple paralellized Boltz-2 workers. B) Generative Virtual Screens: We make use of a molecular generation model to sample batches of compounds to be annotated by Boltz-2. The compounds gradually build a sampled annotated library which is dynamically used to asynchronously train the generative model and generate additional compounds.

**Figure 10:**
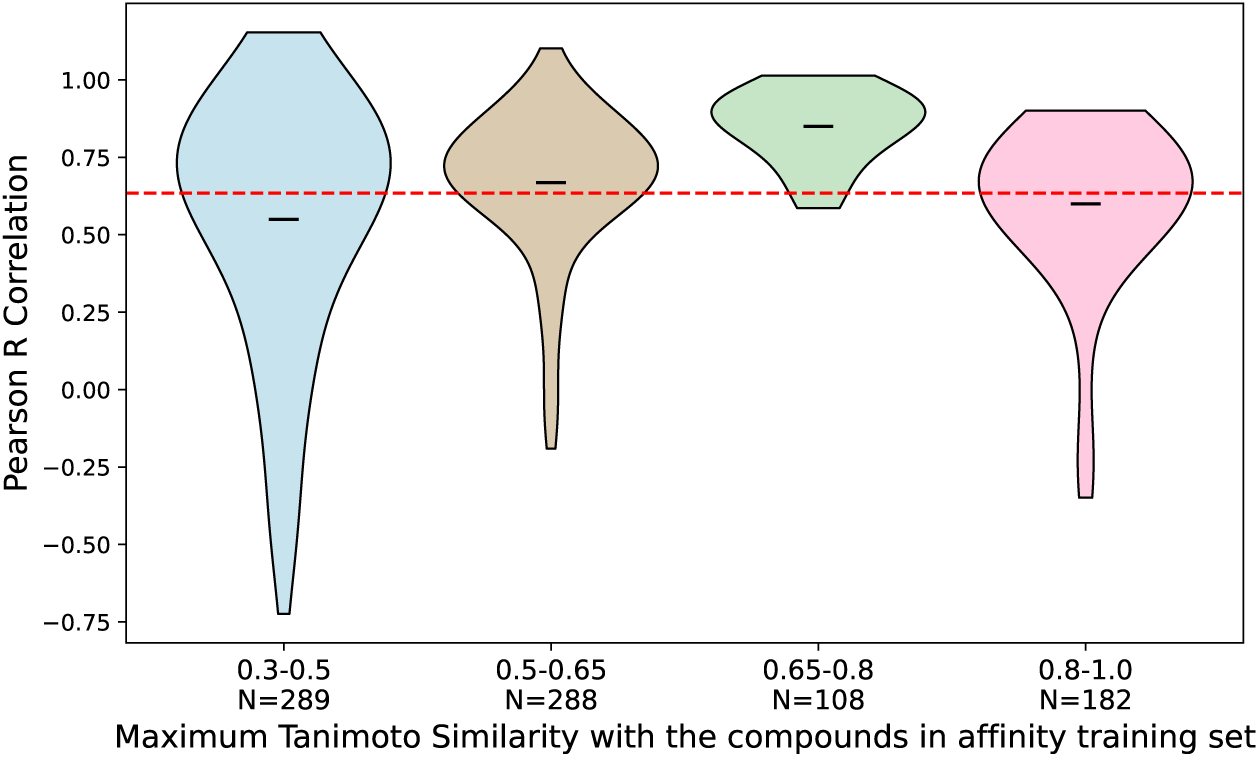
Relationship between compound similarity and prediction performance on the FEP+ benchmark. For each test compound, we compute the maximum Tanimoto similarity to any compound in the affinity training set. We then aggregate these values by computing the per-assay mean of the maximum similarity scores, and group the resulting assays into similarity bins: [0.3, 0.5], [0.5, 0.65], [0.65, 0.8], and [0.8, 1.0]. Each violin plot shows the distribution of per-assay Pearson correlation coefficients between predicted and ground truth affinities. The dashed red line indicates the weighted average of the per-assay Pearson correlations. We observe no strong dependence between compound similarity and predictive performance.

We further extended this screening pipeline using a generative approach. Boltz-2 was coupled with SynFlowNet [Cretu et al., 2024], a GFlowNet-based molecular generator designed to sample molecules from Enamine’s 76B REAL space (details in Appendices B.6 and C.3). This generative screen offers a scalable alternative to fixed libraries by exploring synthesizable chemical space beyond off-the-shelf compounds. After scoring, filtering, and diversity selection, we submitted 10 de novo candidates for ABFE simulation (see Appendix E.3). All selected compounds from the SynFlowNet stream are predicted to bind TYK2, with higher affinity on average compared to fixed screens, and while requiring substantially less computational budget than the HLL screen (117k Boltz-2 evaluations for SynFlowNet against 460k evaluations for HLL). Visualisations of all the selected compounds and of the top-2 ABFE-scored ligand-protein complexes for each stream are presented in Appendix E.3.3. Finally, in Appendix E.3.4, we further assess the novelty of the SynFlowNet-generated compounds by examining their Tanimoto similarity with known binders from the PDB that are part of the structure module training data, and find that the generated compounds do not exhibit significant similarity to public TYK2 binders. We note that these results might be optimistic given that Boltz-2 performs well on this target based on the protein-ligand benchmark data [Hahn et al., 2022], achieving Pearson R = 0.83.

Together, these results demonstrate how Boltz-2 enables structure-based prioritization at a large scale. By addressing both performance and scalability, Boltz-2 expands the scope of target-based in-silico optimization to large scale, encompassing hit discovery, hit-to-lead, and lead optimization.

## 6 Limitations

Despite the progress made in this work for structure and binding affinity prediction, we acknowledge several remaining limitations of the model that we aim to address in future work.

### Molecular Dynamics

While there are clear improvements over Boltz-1, the model does not significantly deviate from other baselines such as AlphaFlow or BioEmu. The current model used a relatively small MD dataset at the later stages of training, with minor architectural changes to account for multiple conformations. Additional changes from the modeling perspective, as well as the datasets used, are required to further improve its capabilities.

### Remaining challenges for structure prediction

While we see a consistent improvement in the structure prediction performance from Boltz-1 to Boltz-2, the model does not significantly deviate from the structure prediction performance of its predecessors. This similarity is primarily due to the use of largely identical structural training data, a similar architectural design, and withstanding limitations in predicting complex interactions, particularly within large complexes. In addition, the model still often fails to capture large conformational changes, such as those that can be induced by binding.

### Accurate structures for affinity predictions

Boltz-2 relies on predicted 3D protein–ligand structures and reliable trunk features as input to the affinity module. If the model fails to identify the correct pocket or inaccurately reconstructs the binding interface or conformational state of the protein, down-stream affinity predictions are unlikely to be reliable. This is particularly relevant in biological contexts where cofactors are essential for binding, given that in its current form, the affinity module does not explicitly handle such cofactors, including ions, water, or multimeric binding partners. Finally, an insufficiently large affinity crop size could be limiting if important long-range interactions are truncated or if the crop does not include the corresponding pocket for each binder, e.g., in the case of both orthosteric and allosteric modulators.

### Understanding the range of applicability of the affinity module

Despite the progress on affinity predictions, we notice in Figures 12-14 that the performance varies strongly between assays. Further work is needed to determine the source of this variance in performance, whether it stems from, e.g., inaccuracies in predicted structures, limited generalization to distinct protein families, or insufficient robustness to out-of-distribution small molecules.

**Figure 11:**
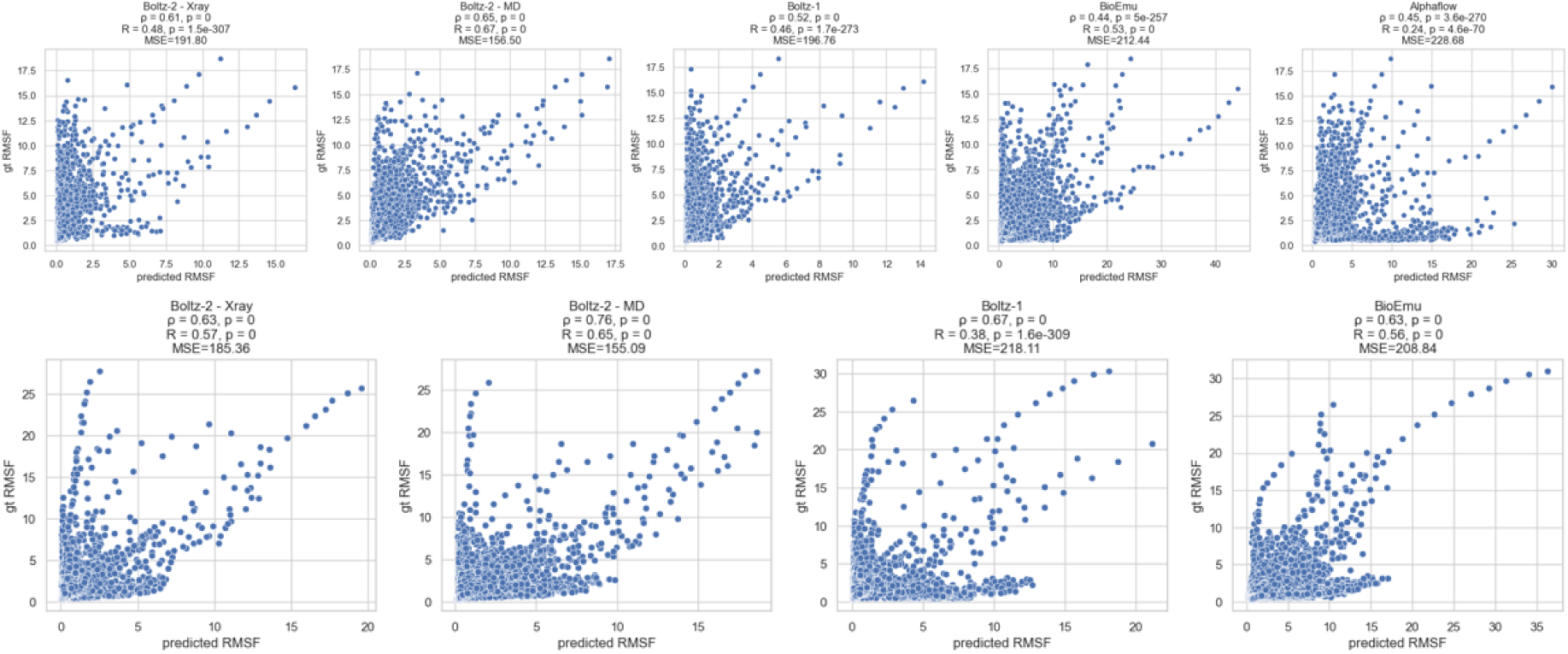
Global RMSF Spearman, Pearson and MSE metrics for the mdCATH (top) and ATLAS (bottom) holdout sets.

**Figure 12:**
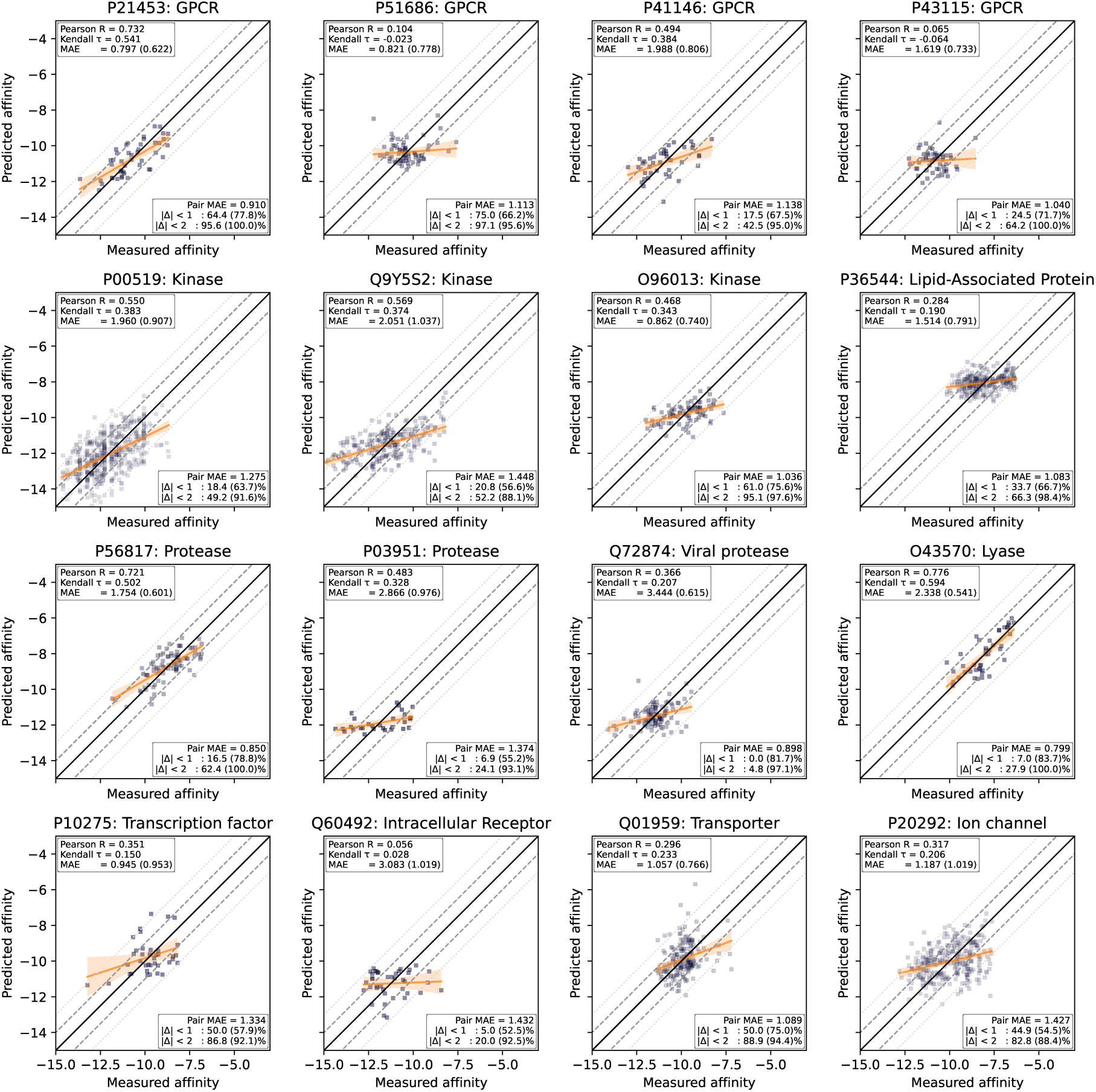
Assay in the validation set, in kcal/mol. The grey lines represent an absolute error of 1 and 2 kcal/mol. |Δ| *< ɛ* represents the percentage of the data that falls within an absolute error of *ɛ*. The number in parenthesis represent the centered metrics.

**Figure 13:**
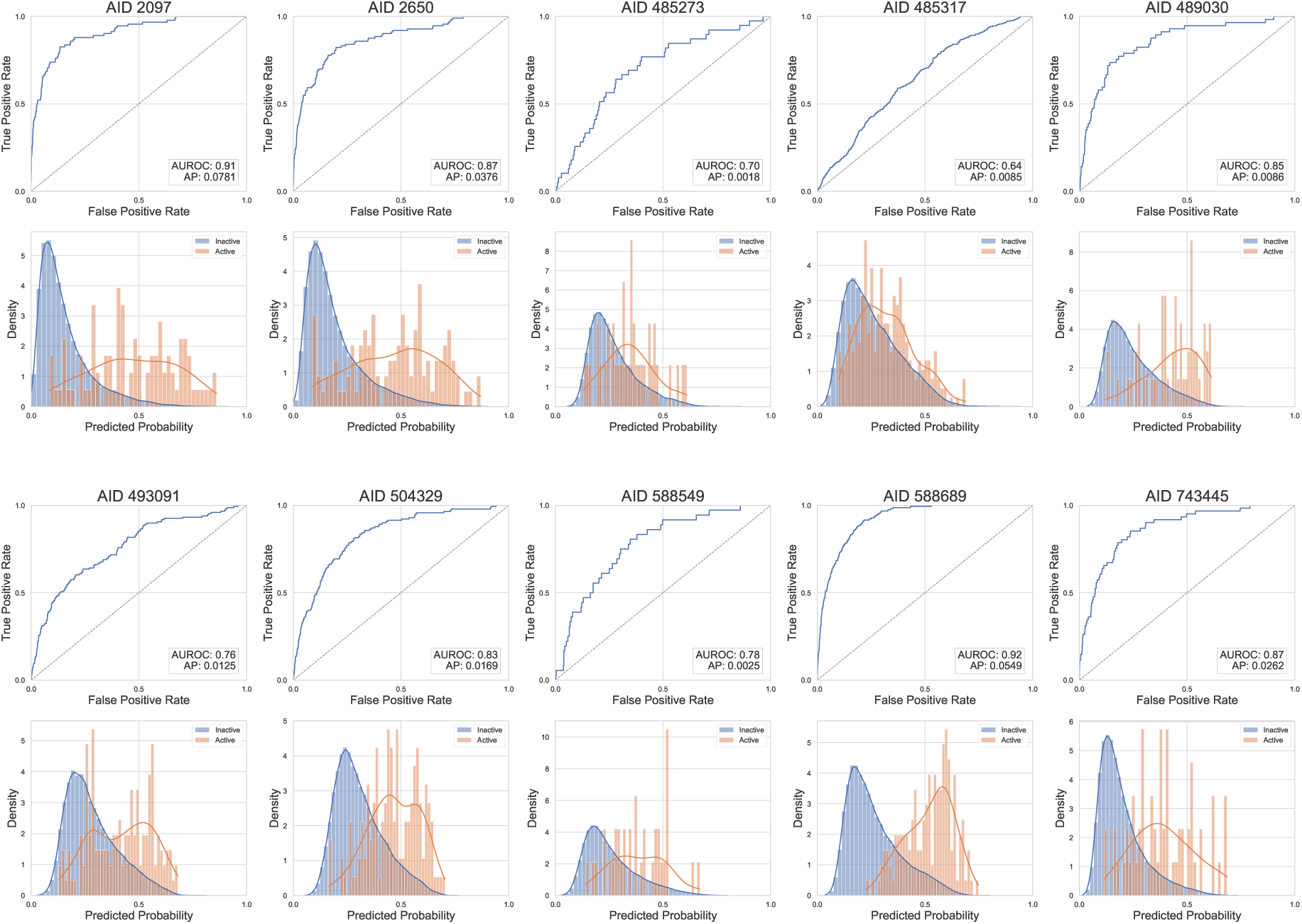
Receiver operating characteristic curves and histograms of binder and decoy distributions for the assays in the MF-PCBA test set.

**Figure 14:**
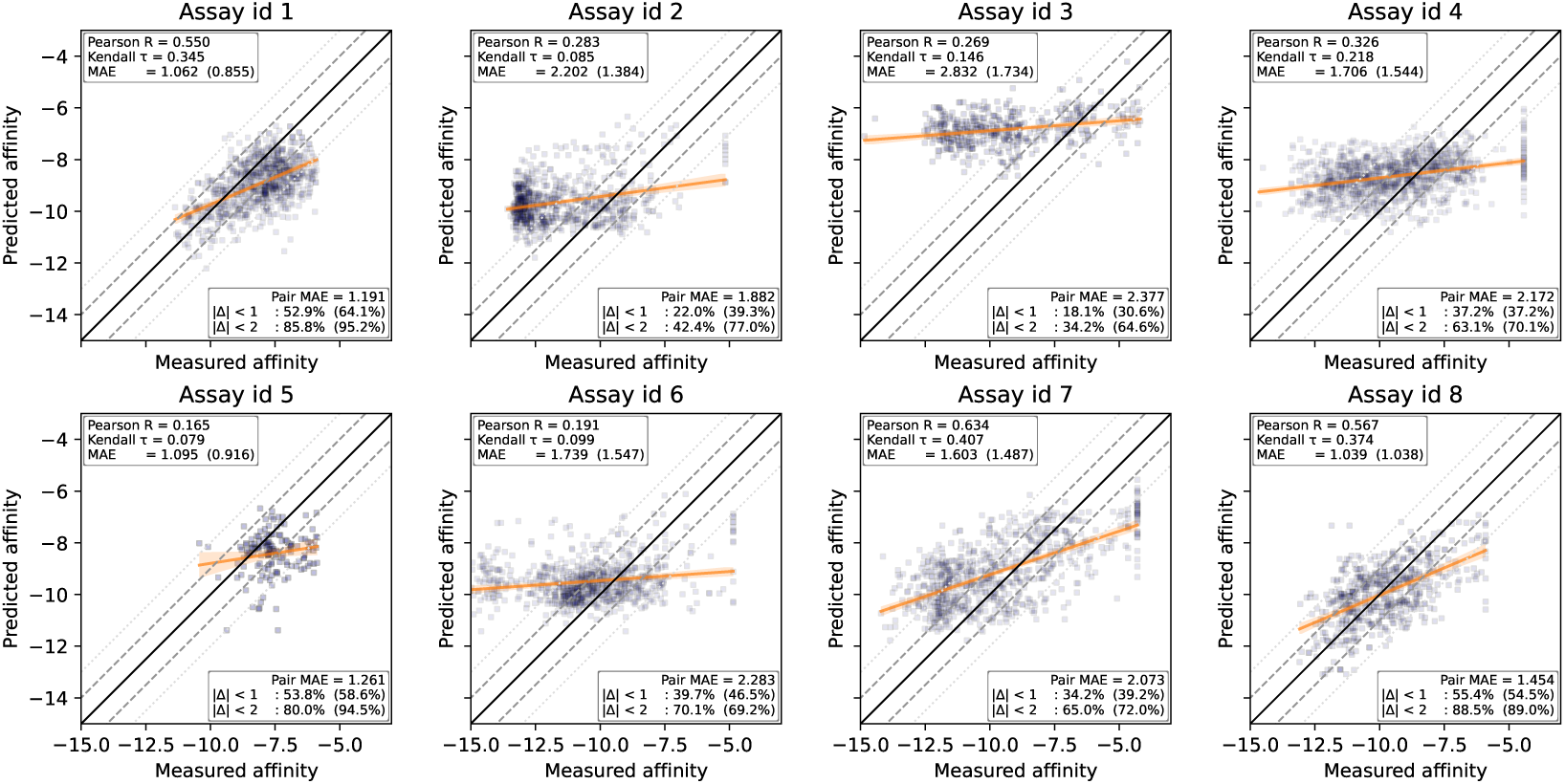
Boltz-2 predictions on 8 blinded targets from private assay, in kcal/mol. The grey lines represent an absolute error of 1 and 2 kcal/mol. |Δ| *< ɛ* represents the percentage of the data that falls within an absolute error of *ɛ*. The number in parenthesis represent the centered metrics.

## 7 Conclusion

We introduce Boltz-2, a new structural biology foundation model that advances the frontiers of both structure and affinity prediction. Boltz-2 builds on the co-folding capabilities of its predecessor with improved physical plausibility, fine-grained controllability, and a better understanding of local dynamics. Our results show that Boltz-2 performs competitively across a broad range of structure prediction tasks, including challenging modalities and conformational ensembles derived from MD. Crucially, Boltz-2 is, to our knowledge, the first AI model to approach the accuracy of FEP methods for predicting binding affinities on the FEP+ benchmark, while offering orders-of-magnitude gains in computational efficiency. For affinity values, Boltz-2 demonstrates strong retrospective and prospective performance in both hit discovery, hit-to-lead and lead optimization settings, as observed on many assays across public benchmarks, private benchmarks, and virtual screening workflows. Coupled with a generative model for small molecules, Boltz-2 enables an end-to-end framework for de novo binder generation, which is validated through ABFE simulations on the TYK2 protein. Despite these advances, several limitations remain, as outlined above. Addressing these issues will require future work in expanding and curating training data, refining model architecture, and integrating additional biochemical context.

By releasing Boltz-2 and its training pipeline under a permissive license, we aim to support the growing community working at the intersection of AI and molecular science. We hope Boltz-2 will serve as a foundation for further advances in drug discovery, protein design, and synthetic biology, expanding the boundaries of what is computationally possible in biomolecular modeling.

## Acknowledgment

We would like to thank Tim O’Donnell, Richard Qi, Ji Won Kim, Sergey Ovchinnikov, Rachel Wu, Felix Faltings, Kyle Swanson, Andreas Krause, Matteo Aldeghi, Francesco Capponi, Demitri Nava, Dylan Reid, Miruna Cretu, and Liam Atkinson for invaluable discussions and feedback around this work. We are grateful to all the members of the Boltz community, many of whom have contributed fixes, features, or helpful feedback that have helped improve the public repository and this project.

We would like to thank Recursion’s High Performance Computing team for their work in establishing, maintaining, solving, and optimizing the GPU usage through the duration of the project, most notably Caden Ellis and Joshua Fryer. We would like to thank Recursion’s physics-based team for helping evaluate poses generated by the model and benchmark the affinity against physical baselines such as docking and FEP, notably Geoff Wood, Gail Bartlett, Arnaldo Filho, Richard Bradshaw, Zhiyi Wu, as well as Thomas Grigg and Oliver Scott for helping with purchasability validation checks. We would like to thank the Valence Labs team for helping us with feedback on the data curation, model training, model evaluation, medicinal chemistry and for reading and proofing the paper, notably Austin Tripp, Michel Moreau, Cristian Gabellini, Lu Zhu, Prudencio Tossou, and Emmanuel Noutahi.

Part of the GPU resources necessary to complete the project were provided by National Energy Research Scientific Computing Center (NERSC), a Department of Energy Office of Science User Facility, via NERSC award GenAI@NERSC. The team from MIT was also supported by the Abdul Latif Jameel Clinic for Machine Learning in Health, the NSF Expeditions grant (award 1918839: Collaborative Research: Understanding the World Through Code), the DTRA Discovery of Medical Counter-measures Against New and Emerging (DOMANE) Threats program, the MATCHMAKERS project supported by the Cancer Grand Challenges partnership (funded by Cancer Research UK (CGCATF-2023/100001), the National Cancer Institute (OT2CA297463) and The Mark Foundation for Cancer Research), the Machine Learning for Pharmaceutical Discovery and Synthesis (MLPDS) consortium, the Centurion Foundation and the BT Charitable Foundation.

## Supplementary Material

### A Data

#### A.1 Structural Data

The training data for Boltz-2 can be divided into two categories, structural and affinity.

The data used for the structural training is an extension of the Boltz-1 data from the PDB [Berman et al., 2000] to include the extractions of ensembles and B-factors [Sun et al., 2019] for supervision and templates for training. Beyond the PDB, it further integrates datasets obtained from molecular dynamics simulations as well as distillation datasets generated with AlphaFold2 and Boltz-1 predictions.

##### A.1.1 PDB data

We process every structure in the PDB following a pipeline similar to those previously described in Boltz-1 [Wohlwend et al., 2025] and AlphaFold3 [Abramson et al., 2024]:

- We use every PDB structure up to the training date cutoff of 06/01/2023. We parse the Biological Assembly 1 from these structures.
- For each polymer chain, we use the reference sequence and align it to the residues available in the structure.
- For ligands, we refer to the CCD dictionary to get the reference ligand and atom composition. We compute up to 10 3D conformers per ligand and sample one at random during training.
- When multiple models are present in the file, such as for NMR structures, we individually process each frame, ensuring that the atomic composition is consistent across frames for the full complex.
- We remove large complexes that are over 7MB or with more than 5000 residues.
- We apply the same filters as AlphaFold3, namely excluding crystallization aids and other non-biologically relevant ligands, removing clashing chains, and filtering out chains with fewer than 4 resolved residues or composed only of unknown residues.
- We compute multiple-sequence alignments for every protein chain (and only protein chains) using ColabFold search. Once monomeric MSAs are produced, we assign a taxonomy ID to every sequence in every MSA using their Uniref100 IDs as reference, if any. The preprocessing of the MSAs is analogous to AlphaFold3.
- We produce template hits for protein chains as described in AlphaFold3, using hmmbuild and hmmsearch on PDB sequences deposited at least 60 days prior to any given query’s deposition date.
- We extract, when available, the B-factors from PDB entries.

##### A.1.2 Molecular Dynamics data

Multiple publicly available molecular dynamics datasets were used for training:

1. *MISATO* : We downloaded the MD dataset from Siebenmorgen et al. [2024], containing NVT trajectories simulated at 300K for 8ns. Trajectories with multi-residue ligands (such as glycans) or modified peptides as ligands are discarded. Ligand matchings to CCD codes are performed by using the reference and mol2 files from PDBbind [Liu et al., 2015]. Trajectories where the ligand floats away from the protein for more than 12^°^A at any frame are removed. Structures in the dataset may contain chain breaks that differ from the corresponding original entries in the PDB. To match these fragments to the appropriate chains and entities in the PDB, their sequences are matched to both the reference author sequence and the structure-derived sequences in the biological assembly 1 with Align.PairwiseAligner. Entries with less than 90% similarity are discarded. Non-overlapping chains that match substrings of the original PDB sequences are greedily merged when possible. Molecular types, entity information and MSA information are assigned after matching to the corresponding parsed PDB data. Polymer and ligands are then parsed with the same pipeline used to parse the PDB, as described above. All 100 frames from the 8ns are parsed and used for training. After all filters are applied, the dataset results in 11,235 systems.
2. *ATLAS* : We downloaded the MD dataset from Vander Meersche et al. [2024], containing NPT trajectories simulated at 300K for 100ns. Structures are matched to their corresponding PDB entries. Replica trajectories are aggregated, and 100 frames are uniformly sampled at random from the final 10 nanoseconds of each trajectory for training, resulting in a final dataset of 1,284 proteins.
3. *mdCATH* : Trajectories from Mirarchi et al. [2024] containing NVT trajectories at 320K with varying simulation times up to 500ns were used. For training, only the last 10% of the trajectory was utilized. The final dataset comprises 5,270 systems.

##### A.1.3 Distillation data

For training the Boltz-2 model, we used several datasets generated by distilling predictions from the Boltz-1 model, while applying appropriate filters. For all datasets generated from Boltz-1, we employed 3 recycling steps in the trunk and generated 3 diffusion samples per example.

1. *RNA distillation*: Following AlphaFold3, we clustered Rfam (v14.9) [Kalvari et al., 2021] using MMSeqs2 [Steinegger and Söding, 2017] with 90% identity and 80% coverage. To form the distillation set, Boltz-1 predictions for cluster representatives are filtered to those where the maximum average predicted distance error (PDE) ≤ 2.0.
2. *Protein-DNA distillation*: Our Protein-DNA distillation data is constructed similarly to AlphaFold3. Using the JASPAR 2024 release (specifically, the CORE collection), we first find transcription factor profiles with matching gene IDs across two high-throughput SELEX datasets [Jolma et al., 2015, Yin et al., 2017]. For each filtered profile, a protein sequence is assigned in two ways: i) using the canonical protein sequence under the profile’s Uniprot ID and ii) searching for the sequence in the two SELEX datasets (with matching gene ID) with the highest similarity to the Uniprot sequence. Sequence similarity is calculated using KAlign v2.0, computed as the number of non-gap matches between the two sequences divided by the minimum length of pre-aligned sequences. Unlike AlphaFold3, we did not apply any sequence clustering. To generate binding DNA sequences for each protein sequence, we use the corresponding JASPAR profile’s position frequency matrix (PFM) to sample 10 single-stranded motifs. For each distillation example, the inputs include the protein sequence, the single-strand DNA sequence and its corresponding reverse complement. After generating Boltz-1 predictions, we filtered examples to those that satisfied all the following conditions PDE ≤ 2.0, maximum interface predicted distance error (iPDE) ≤ 1.0 and minimum interface predicted TM-Score (ipTM) ≥ 0.7.
3. *RNA-Ligand distillation*: We use the R-SIM [Krishnan et al., 2023] dataset comprising ∼ 2500 examples of RNA-small molecule affinities. Predictions from Boltz-1 for these examples are filtered to those with either iPDE ≤ 1.0 or (ipTM) ≥ 0.7.
4. *Protein-Ligand distillation*: We construct a dataset of protein-ligand distillation from BindingDB and ChEMBL that were excluded from the main hit-to-lead affinity training set due to having too few compounds in the assay. The full parsing and filtering criteria applied are described in Appendix A.2. The distillation set was formed by filtering Boltz-1 predictions to examples with a maximum interface predicted distance error (iPDE) ≤ 1.0 and a minimum interface predicted TM-Score (ipTM) ≥ 0.9.
5. *TCR-pMHC distillation*: We generate the TCR self-distillation dataset (class I and class II) from VDJdb [Goncharov et al., 2022]. We retain only paired TCR entries with complete gene annotations, enabling accurate reconstruction of full-length TCR sequences using Thimble, the bulk-processing version of Stitchr [Heather et al., 2022]. Entries flagged with reconstruction errors by Thimble are removed. The inputs for Boltz-1 contained TCR alpha and beta sequences (cropped to 9 residues beyond the CDR3 region), peptide sequences, and MHC alpha/beta sequences (cropped at 180 residues). Multiple Sequence Alignments (MSAs) are generated using ColabFold [Mirdita et al., 2022] and included for TCR and MHC chains, but not peptides. The distillation set was formed by filtering Boltz-1 predictions to examples with a (iPDE) ≤ 1.0, PDE ≤ 1.0, and (ipTM) ≥ 0.8.
6. *MHC-I and MHC-II* : We generate the pMHC self-distillation dataset (class I and class II) from the Immune Epitope Database (IEDB) [Vita et al., 2025]. We initially filter for epitopes with clearly annotated MHC alleles, remove mutant epitopes, and exclude sequences already present in the Protein Data Bank (PDB). Class I epitopes were restricted to 8-12 residues and class II epitopes to 13-25 residues. For class 1, we included MHCs from across multiple species. For class 2 MHCs, we only use human MHC alleles and discard entries for which we could not determine standardized and paired MHC sequences. For each MHC allele, we crop sequences to a maximum length of 180 residues. To limit redundancy, we sample up to 100 sequences per allele for class I and up to 200 per allele pair for class II. The distillation set is formed by filtering Boltz-1 predictions to examples with a (iPDE) ≤ 1.0, PDE ≤ 1.0, and (ipTM) ≥ 0.85.
7. *AlphaFold Database (AFDB) distillation:* In order to construct a protein monomer distillation set, we begin with uniref30 and find the overlap between those sequences and the uniclust multiple sequence alignments provided by OpenFold. We then fetch structures from the AFDB where we impose a minimum global lDDT of 0.5. This procedure results in a monomer distillation of about 5 million proteins.

##### A.1.4 Training sampling weights

Table 2 summarizes the various datasets used for the structure training and details their relative sampling weights. As shown, from the PDB we upsample interfaces containing antibodies and TCR, as these are specific modalities that we wanted the model to improve on, as well as proteins similar to SARS-CoV2-Mpro as we had planned to participate in the Polaris-ASAP competition.

**Table 2:** Breakdown of training data sources and their relative sampling weights. Sampling clusters indicate how elements are counted in the sampler that selects the complexes to train on and also determines where the first token of the crop is placed.

| Dataset | Source | Sampling Clusters | Sampling weight |
| --- | --- | --- | --- |
| PDB | experimental | chains & interfaces | 0.455 |
| Antibodies upsampling | experimental | interfaces | 0.025 |
| TCR upsampling | experimental | interfaces | 0.004 |
| SARS-Cov2-Mpro upsampling | experimental | interfaces | 0.015 |
| AFDB | AF2 distillation | chains | 0.380 |
| Protein-ligand | Boltz-1 distillation | interfaces | 0.031 |
| RNA | Boltz-1 distillation | chains | 0.045 |
| DNA-protein | Boltz-1 distillation | interfaces | 0.020 |
| RNA-ligand | Boltz-1 distillation | interfaces | 0.001 |
| TCR | Boltz-1 distillation | interfaces | 0.012 |
| pMHC-I | Boltz-1 distillation | interfaces | 0.006 |
| pMHC-II | Boltz-1 distillation | interfaces | 0.006 |
| ATLAS | MD | chains | 0.003 |
| MISATO | MD | chains & interfaces | 0.230 |
| mdCATH | MD | chains | 0.010 |

##### A.1.5 Validation dataset

###### PDB data

Our training, validation and test splitting strategy largely follows Boltz-1 procedure [Wohlwend et al., 2025]. We first cluster the protein sequences in PDB by sequence identity with the command mmseqs easy-cluster … –min-seq-id 0.4 [Hauser et al., 2016]. Then, we select all structures in PDB satisfying the following filters:

1. Initial release date is before 2023-06-01 (exclusive) and 2024-01-01 (inclusive).
2. Resolution is below 4.5 ^°^A.
3. All the protein sequences of the chains are not present in any training set clusters (i.e. before 2023-06-01).
4. Either:

a. No small-molecule is present.
b. At least one of the small-molecules exhibits a Tanimoto similarity of 0.85 or less to any small-molecule in the training set. Here, a small-molecule is defined as any non-polymer entity containing more than one heavy atom and not included in the ligand exclusion list.
c. The small-molecule satisfies Lipinski’s Rule of Five.

We further refine through the following steps:

1. Retain structures with at most 1024 residues.
2. Exclude complexes with more than 20 entities.
3. Retaining all the structures containing RNA or DNA entities.
4. Iteratively adding structures containing small-molecules or ions under the condition that all their protein chains belong to new unseen clusters.
5. Iteratively adding multimeric structures under the condition that all the protein chains belong to new unseen clusters. These are further filtered by randomly keeping only 90% of the passing structures.
6. Iteratively adding monomers under the condition that their chain belongs to a new unseen cluster. These are further randomly filtered out by keeping only 60% of the passing structures.

This results in a total of 398 structures from PDB in our validation set.

###### MD data

Given that all entries from the MD datasets correspond to PDB structures released before the validation cutoff date of 2023-06-01, we constructed test sets by greedily adding structures with the smallest number of members within their cluster group. Here, we use the same clusters defined in the general PDB dataset with a sequence identity threshold of 0.4. Trajectories from the selected clusters and all of their cluster members are then removed from the training set across all 3 MD datasets. This procedure is repeated until we achieve a desired test set of 40 complexes per dataset.

#### A.2 Binding Affinity Data

We construct a large-scale dataset for training and evaluating our model, comprising both continuous affinity measurements (e.g., *K_i_*, *K_d_*, *IC*_50_, *AC*_50_, *EC*_50_, *XC*_50_) and binary labels (binder vs. decoy). The dataset integrates information from a diverse set of public sources:

- **PubChem (1.8.1):** A large public repository maintained by the NIH that contains bioactivity data across a broad range of targets and compounds [Kim et al., 2023].
- **ChEMBL (v34):** A manually curated database of bioactive molecules with drug-like properties, containing standardized binding and functional assay data [Zdrazil et al., 2024].
- **BindingDB:** A public database of measured binding affinities for protein–ligand interactions, with a strong emphasis on *K_i_* and *K_d_* values from medicinal chemistry studies [Liu et al., 2007].
- **CeMM Fragment Dataset:** A dataset of fragment-screening results containing validated binders from fragment-based drug discovery campaigns [Offensperger et al., 2024].
- **MIDAS Metabolite Data (University of Utah):** A set of small-molecule metabolites tested in biochemical binding assays by the University of Utah [Hicks et al., 2023].

Since our model leverages protein–ligand complex structures as input, it is essential to ensure the structural quality of the training data. To this end, we apply a filtering strategy at the assay level that avoids introducing selection bias. For each assay, we compute the average predicted interface pTM (ipTM) score across a set of 10 random binders using the Boltz-2 confidence module. Assays are retained only if this average exceeds 0.75, ensuring that the dataset contains structurally reliable examples suitable for structure-based learning.

We use the ChEMBL Structure Pipeline [Bento et al., 2020] to standardize all ligand molecules across the datasets. This pipeline applies a series of cheminformatics preprocessing steps designed to ensure consistency. The resulting standardized SMILES are then used as input to Boltz-2.

##### A.2.1 Affinity values

We curate continuous affinity data primarily from ChEMBL and BindingDB, using the following filtering and standardization steps:

###### ChEMBL data extraction

We extract binding measurements from ChEMBL using the following criteria:

- Filter to confidence score equal 9 (maximum confidence) to retain high-quality structure–activity annotations.
- Target type restricted to SINGLE PROTEIN to train on high-quality structures.
- Filter to biochemical or functional assays.
- Filter to affinity measurements with standard type in {’Ki’, ’Kd’, ’IC50’, ’XC50’, ’EC50’, ’AC50’}.
- Exclude sources flagged as unreliable.
- Parse and retain protein mutation annotations when available.
- Store both the assay ID and activity qualifier for downstream processing.

###### BindingDB data extraction

We retain only records not already covered by ChEMBL, using the following protocol:

- Use the BindingDB DOI identifier as the assay ID.
- Exclude proteins that report more than 1 chain.
- Retain associated activity qualifiers.
- Parse the protein sequence reported by BindingDB.

###### General curation

Across both datasets, we:

- Remove PAINS (Pan-Assay Interference Compounds) to eliminate molecules known to produce assay artifacts or interfere with diverse biochemical readouts.
- Filter out molecules with more than 50 heavy atoms.
- Partition the curated affinity data into two subsets: 1) hit-to-lead affinity values, representing optimization-stage datasets, and 2) hit affinity values representing earlier-stage binding screens. This separation allows us to handle affinity values with > qualifiers appropriately: in hit discovery, they are treated as decoys, while in hit-to-lead, they are interpreted as censored measurements, reflecting the uncertainty of whether the compound is a weak binder or a true non-binder.

###### Hit-to-lead curation

For optimization-stage assays, we apply stricter filters to minimize the noise from these assays:

- Convert all affinity values to logarithmic scale with 1 *µ*M as the reference unit.
- Remove assays with low average pairwise Tanimoto similarity (*<* 0.25), retaining those aligned with hit-to-lead settings where actives are structurally related.
- Exclude assays with fewer than 10 data points, to further focus the model on fitting affinity differences of similar molecules and not global trends.
- Discard assays with low activity standard deviation (*<* 0.25), as they do not help to understand activity cliffs.
- Exclude assays with fewer than 10 unique activity values or where the unique-to-total ratio is less than 0.2, as these are likely to come from low-accuracy assays.
- Discard data with qualifiers <.
- Discard data with qualifiers > and activity value *<* 10 µM.
- Remove assays containing any activity value *<* 10^−6^ *µ*M, as these often indicate incorrectly reported units or annotation errors.

###### Hit-discovery curation

For screening assays:

- Retain only assays with at least 100 data points.
- Retain chemically diverse assays (average pairwise Tanimoto similarity *<* 0.25), discarding hit-to-lead assays with low diversity.
- Label as *inactive* all entries with *>* qualifiers, as *active* those with = and affinity *<* 2.0 µM and discard everything else.

Despite our effort, our affinity value curation only scratches the surface of what is possible for constructing high-quality training datasets. Future work could pursue more rigorous standardization by, for instance: (1) applying the Cheng–Prusoff equation to convert inhibition assay values (e.g., *IC*_50_) into *K_i_* estimates for more direct comparability; (2) performing deeper assay-level vetting to exclude data from cell-based assays, low-purity protein preparations, or other sources known to introduce noise; and (3) further removing artifacts and confounding signals through advanced filtering or metadata-driven heuristics. However, such refinement is very challenging in practice, as assay metadata are often inconsistently reported, difficult to parse, and require close collaboration with domain experts deeply familiar with the biological and experimental nuances of each assay.

##### A.2.2 Binary labels

Binary classification data are derived primarily from HTS assays in PubChem and supplemented with fragment, metabolite binding data, and a synthetically generated set of decoys.

###### PubChem HTS curation

We apply the following filters to construct a reliable binary dataset:

- Retain only assays with at least 100 tested compounds.
- Retain assays with a hit rate (actives/total) *<* 0.1.
- For each (protein sequence, SMILES) pair, we query PubChem for matching entries that report an affinity value measurement and are explicitly labeled as Active. Only compounds meeting both criteria are retained; all others are discarded. Through cross-referencing with available confirmatory (secondary) screens, we estimate that approximately 40% of the compounds labeled as actives in high-throughput primary screens may be false positives.
- Remove PAINS compounds.
- Subsample the decoy set to achieve an approximate 1:9 ratio of binders to decoys per assay.

###### CeMM Fragment

We apply the following filters to construct a reliable dataset:

- Remove all fragments labeled with low confidence (score = 1).
- Label as binders all fragments with medium or high confidence (scores = 2 or 3).
- Label as inactives all fragments explicitly marked as decoys (score = 0).
- Subsample the decoy set to achieve an approximate 1:9 ratio of binders to decoys per assay.

###### Synthetic decoys

To construct a reliable set of synthetic decoys for binary classification, we apply the following procedure:

- Each hit-to-lead compound with an experimentally measured affinity is paired with a single decoy molecule.
- Decoys are sampled from the pool of hit-to-lead molecules to ensure distributional consistency between binders and decoys. This avoids trivial shortcuts where the model could distinguish actives from decoys based solely on distributional differences, rather than true binding signal.
- Assuming that hit-to-lead compounds are selective, we minimize the likelihood of false negatives by sampling decoys from molecules with a Tanimoto similarity *<* 0.3 to any known binder of targets belonging to the same 90% sequence identity cluster as the current target. This constraint reduces the chance of accidentally including active compounds as decoys.

Binary label curation poses unique challenges, particularly when integrating data from high-throughput screening (HTS). Although HTS datasets are valuable due to their scale and realistic chemical diversity, they are susceptible to multiple sources of systematic noise and artifacts. Examples include false positives arising from promiscuous binding (e.g., PAINS compounds or colloidal aggregators), interference artifacts such as luciferase inhibition or fluorescent signal quenching, and biological noise introduced through cell-based assays where the measured signal may not directly reflect binding to the intended protein target. Our current strategy—matching binary labels to corresponding affinity measurements—provides an initial filter but does not fully ensure reliability or biological relevance. For instance, matched affinity data often stems from secondary assays whose quality or target specificity we have not systematically validated. Achieving a robust binary curation would require deeper metadata interrogation to confirm assay quality, target specificity, and orthogonal validation outcomes. However, inconsistent metadata annotation and reporting across public repositories substantially complicate such efforts, necessitating close collaboration with experimentalists and domain specialists who possess detailed knowledge of assay methodologies and underlying biology.

### B Model

#### B.1 Tokenization and Featurization

Shared across all modules are the tokenization of the biomolecular complexes and the featurization of each atom and each token.

##### Tokenization

We use the following tokenization scheme: Every protein is tokenized at the amino-acid level, every RNA and DNA at the nucleotide level, and other biomolecules are tokenized at the atomic level. Unlike AlphaFold3, Chai-1, and Boltz-1, where non-canonical amino acids and nucleotides are tokenized at the atomic level, we keep them as a single token as well.

##### Featurization

Compared to Boltz-1, Boltz-2 has the following additional features that are given as input to the model. At the single token level, a cyclic flag distinguishing cyclic polymers from acyclic ones, a modified flag distinguishing non-canonical amino acids or nucleotides, a one-hot encoding for different experimental method types, and a molecular type feature encoding whether a token belongs to a protein, DNA, RNA, or other. At the pairwise token level, a bond type feature distinguishes the order/aromaticity of bonds between pairs of tokens. Moreover, the relative positional encoder was modified to have cyclic-offset positional encodings for cyclic polymers (similar to Rettie et al. [2025]) and only have the relative chain encoding between symmetric chains (similar to ProteinX [Chen et al., 2025]). Finally, for sequences in the MSA, we add an additional binary feature to every token representing whether or not it is part of a paired sequence.

#### B.2 Trunk and Denoising Modules Architecture

##### Trunk Module

At high-level, the architecture of the model in the trunk is similar to that of Boltz-1 with a few exceptions:

1. Boltz-2 utilizes a template module similar to AlphaFold3.
2. The number of PairFormer layers are increased from 48 to 64.
3. For a majority of the trunk, we employed mixed-precision training (using bfloat16) and trifast kernels for triangular attention operations. This allowed us to scale the crop size at training time from 512 (in Boltz-1) to 768 (similar to AlphaFold3).

##### Denoising Module

We used the same denoising module as Boltz-1. The denoising module was trained in float32 precision due to instabilities observed at lower precision.

##### Physical quality

Deep learning based co-folding models such as AlphaFold3, Chai-1 and Boltz-1 suffer from significant physical issues with the poses they generate. These include the presence of chain clashing hallucinations as well as other issues including steric clashes between atoms, slightly incorrect bond lengths and angles, incorrect stereochemistry at chiral centers and stereobonds and aromatic rings predicted to be non-planar. We recently introduced Boltz-steering, a new inference-time technique that, when applied with a set of physics-based potentials on top of Boltz-1 gave rise to Boltz-1x keeping the original geometric accuracy while solving many of these physical issues. Boltz-2 adopts the steering potentials we proposed in Boltz-1x with tuned hyperparameters and a normalization of each potential by the number of elements on which they are applied. Additionally, as presented in the next section, we integrated additional potentials to improve controllability.

#### B.3 Controllability

##### B.3.1 Method conditioning

Boltz-2 is trained with structural data generated with a variety of different experimental methods, including but not limited to: X-ray diffraction, electron microscopy, solution NMR, solid-state NMR, molecular dynamics, distillation from AlphaFold2 and distillation from Boltz-1. As structure prediction methods are reaching experimental accuracy, it is important to teach them to understand the different properties that structures coming from different experimental methods have.

Therefore, at training time, we condition the model to the experimental method to obtain the specific structure by giving its experimental method as input in the single token representations. At inference time, users can decide on which experimental method to use to condition the model’s prediction. As shown in Section 5.2, this conditioning does indeed have an effect on the resulting structure distribution, leading it closer to the one obtained with the desired experimental technique.

##### B.3.2 Templates conditioning and steering

Templates allow the users to feed the model the structures of related biomolecular complexes that might help with the prediction of the complex under analysis. While not very effective when fed with structures within the model’s training set, templates can be particularly useful in settings where users have access to unseen relevant structures or have a strong prior on the complex structure having a particularly similar fold.

While Boltz-1 does not support template conditioning, AlphaFold3 and Chai-1 integrate it. However, they only allow for single chains to be used as templates, and they do not enforce the model to necessarily respect the given template, often leading to no improvements from the template addition. In Boltz-2, we improve on these two fronts: we allow for multimeric templates and we allow the user to strictly enforce that templates are respected via a Boltz-steering potential. Similar to previous models, at this point, we only allow for potentials within protein chains.

###### Template conditioning

We first produce template hits for all monomeric protein chains. During training, we then group the templates by their PDB ID so that if chains A and B yield two templates of the same PDB ID, these will be grouped together and used as a multimeric template. Note that we only do this over protein chains. Following AlphaFold3, we always sample 0 to 4 templates per chain. We do so by keeping a counter for each chain and using template groups as defined above to assign templates to chains with non-zero counters. Whereas previous approaches limit the template mask to only be non-zero along the diagonal (i.e, monomeric templating), in our approach, templates with the same PDB ID are visible to one another during template encoding.

###### Template steering

If the user desires to enforce the potential beyond what the conditioning does, we devised an inference time Boltz-steering potential that pushes the reverse diffusion to place the portion of the structure corresponding to the template (which can be a subset of a chain, such as a pocket, a domain or a loop) to have a structure within *α*_cutoff_ ^°^A of the given template.

For a template with reference atoms *S*_template atoms_, we define the potential as follows.

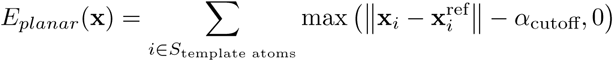

where **x**^ref^ is the position of reference atom *i* after aligning the template to the predicted coordinates.

##### B.3.3 Contacts and pocket conditioning and steering

From experimental data or intuition, structural biologists often have hypotheses about which residues within a complex might interact or which site of a polymer another molecule might bind to. We will define a contact as the specification of a distance constraint between two tokens (residues/atoms), and a pocket as the specification of a distance constraint between one or more tokens and a separate chain/molecule.

Although it did report results on a separately trained model with pocket conditioning, AlphaFold3’s publicly available model does not support any distance constraint specification. Boltz-1 supports the specifications of pocket conditioning, Chai-1 supports both pocket and contact conditioning with flexible distance cutoff. In Boltz-2, we also support both pocket and contact conditioning with flexible distance cutoff. These are added not only like previous models with feature conditioning, in which case the model sometimes takes samples that do not respect these conditions, but also with steering potentials to enforce them.

###### Contact and pocket conditioning

Boltz-1 defined pocket conditioning via features fed into the single token representation. This, however, has the limitation that at most one restraint can be specified. Therefore, in Boltz-2, we feed contact and pocket conditioning as pairwise features between tokens. These features consist of a one-hot encoding of the contact type and an encoding of the distance. The contact type is selected among: no restraint was specified, some were specified but this was not selected, this pair has a pocket-to-binder relationship, this pair has a binder-to-pocket relationship, this pair has a contact relationship (takes precedence). The encoding of the distance *d*, constrained to be 4^°^A ≤ *d* ≤ 20^°^A, is encoded as a concatenation of the normalized distance (*d* − 4)*/*16 and its Fourier embedding with a fixed set of randomly sampled bases.

###### Contact and pocket steering

When defining a contact or a pocket, these can be interpreted as a relationship between two sets of atoms, *S_A_* and *S_B_*, where our goal is to ensure that the smallest distance between the atoms in these sets is less than a threshold *r_AB_*. To do this, we define a time-dependent potential function as follows.

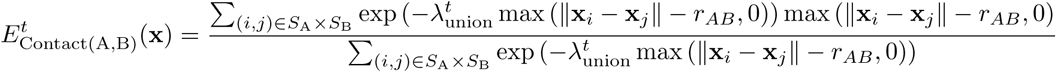

where *λ^t^* is a time-dependent parameter which monotonically increases as t goes from 1 to 0. As *t* → 0, the potential biases the model towards conformers where all pairs of atoms are within a distance of *r_AB_*, and as *t* → ∞, the potential biases the model towards conformers where any pair of atoms is within a distance of *r_AB_* to enable more flexibility in the ligand conformation within the pocket.

#### B.4 Confidence Module Architecture

The confidence module of Boltz-2 has an architecture that resembles that of AlphaFold3’s confidence model. Instead, Boltz-1 has a significantly more expensive confidence model that included a trunk of the same size of the structure prediction trunk (48 layers of PairFormer plus AtomEncoder and MSA modules), which is initialized with the weights of the structure prediction trunk and includes inputs from the DiffusionTransformer final representations. While this larger architecture provides some improvement over the simpler architecture of AlphaFold3 and Boltz-2 confidence model, it comes at a significant cost.

Therefore, we opted for a faster architecture, using eight PairFormer layers (versus the four of AlphaFold3) on top of the final pair token representation of the structure trunk and the encoding of the predicted coordinates. Unlike previous models, we found it beneficial to divide the final heads predicting the PDE and PAE logits into two separate layers, one making the prediction for pairs of tokens within the same chain/molecule and one making the prediction for pairs across different chains/molecules.

#### B.5 Affinity Module Architecture

One of the core challenges in drug discovery is accurately determining whether a small molecule binds to a given protein target and quantifying the strength of this interaction. Boltz-2 enhances this capability through a specialized affinity module designed to address two key prediction tasks:

**Algorithm 1:**
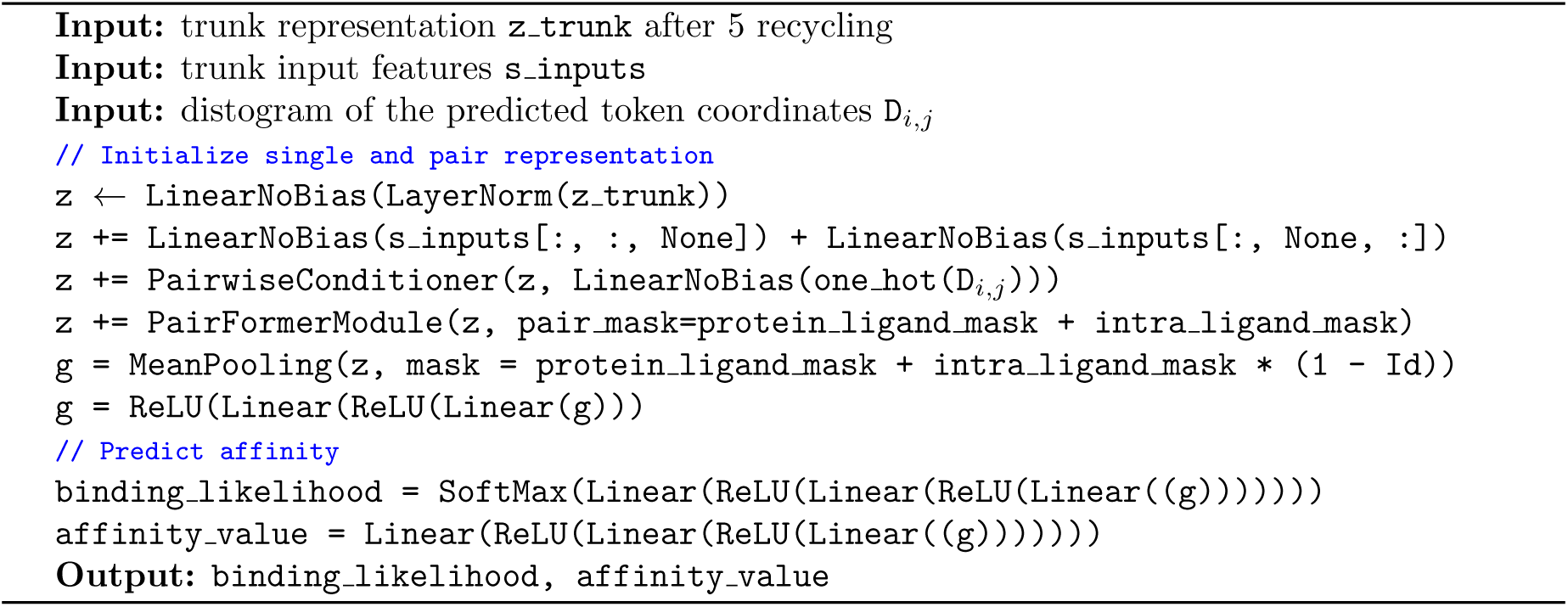
Affinity module

1. **Binding likelihood**: Predicting the likelihood that a small molecule will bind to a specific protein target.
2. **Affinity value**: Quantifying the strength of the interaction between a small molecule and a protein target, measured similar to the half-maximal inhibitory concentration (*IC*_50_).

The binding likelihood head is optimized for identifying potential hits across diverse molecular screening scenarios. In contrast, the affinity value head is specifically tailored to guide hit-to-lead optimization by discerning subtle variations in binding strengths among structurally related molecules targeting the same protein. Algorithm 1 details the full implementation of this affinity module.

##### B.5.1 Affinity Module

The affinity module operates on the structural predictions from Boltz-2, specifically utilizing the input single representations *s*_inputs_ and the final pair representation **z***_ij_* obtained after five recycling iterations. Coordinates fed into the module are selected as the top-ranked structure from five samples generated over 200 diffusion steps each, ranked according to their protein-ligand ipTM-score.

At its core, the affinity architecture comprises a Pairformer model designed to process the interaction pair representations, masking out intra-protein interactions to focus exclusively on protein-ligand interface details. To achieve a scalar representation for the downstream affinity prediction, the module performs mean pooling over all pairwise interactions.

Following pooling, two dedicated multi-layer perceptron (MLP) heads produce distinct outputs: one providing logits for binding likelihood estimation and another regressing continuous affinity values.

##### B.5.2 Affinity Model Ensemble

To improve robustness and overall performance, we train two affinity models with distinct hyperparameters. The models differ in binder-to-decoy loss weighting (*λ*_focal_ = 0.8 vs. 0.6), the number of transformer layers (4 vs. 8), and training duration (one is trained longer while the other is early-stopped). This diversity not only enhances predictive accuracy through ensembling but also serves an important role in downstream molecule generation. When using SynFlowNets for optimization, there is a risk of over-optimizing against a single model’s reward signal. To mitigate this, we use the second model as an independent reference for final filtering, reducing the likelihood of reward hacking and introducing a more stable selection criterion.

We ensemble the models as follows:

- For binary classification, we take the average of the predicted binding likelihoods.
- For affinity regression, we apply a calibrated ensembling strategy. We first compute the mean predicted affinity between models and then apply a molecular weight correction of the form

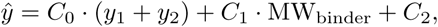

where *y*_1_ and *y*_2_ are the predictions of the two models, *C*_0_, *C*_1_, and *C*_2_ are fitted in the holdout validation set and MW_binder_ is the molecular weight of the binding small molecule.

In Table 3, we report the differences in hyperparameters between the two affinity models used in the ensemble. Both models are trained across 128 A100 GPUs using the AdamW optimizer with a weight decay of 0.001 and a learning rate of 0.0001.

**Table 3:** Extra hyperparameters that differ between the two affinity models trained for the ensemble.

|  | Ensemble member 1 | Ensemble member 2 |
| --- | --- | --- |
| PairFormers layers | 8 | 4 |
| $\lambda_{focal}$ | 0.8 | 0.6 |
| Training samples ($\cdot 10^6$) | 55 | 12.5 |

#### B.6 SynFlowNet Ligand-Generation Model

For SynFlowNet, we use a similar setup to that of Cretu et al. [2024]. The Markov Decision Process (MDP) traversed by the agent represents partial molecules, with the action space comprising both uni-and bi-molecular reactions and of a set of building blocks. The agent sequentially constructs each trajectory by combining these elements. The forward policy *P_F_* is parameterized by a Graph Transformer model [Yun et al., 2019], while the backward policy *P_B_* is a uniform distribution over the backward actions. We employed the trajectory balance loss [Malkin et al., 2022]. All model and training hyperparameters are detailed in Table 4.

**Table 4:** Model and training hyperparameters for our SynFlowNet molecular generation model.

| Hyperparameter | Value |
| --- | --- |
| Numbr of training steps | 11,000 |
| Training batch size | 64 |
| Replay buffer warmup | 500 |
| Maximum trajectory length | 3 |
| Reward function exponent ($\beta$) | 36 |
| Random action probability (exploration) | 0.20 |
| Target policy soft update parameter ($\tau$) | 0.99 |
| Training loss | Trajectory Balance |
| Backward policy $P_B$ type | Uniform |
| Optimizer | Adam |
| Forward policy $P_F$ learning rate | $10^{-4}$ |
| $P_F$ learning rate decay | 2,000 |
| Normalizing constant $Z$ learning rate | $10^{-3}$ |
| $Z$ learning rate decay | 50,000 |
| Graph transformer embedding size | 128 |
| Graph transformer depth | 4 |
| Graph transformer number of heads | 2 |
| Action space reactions (number & source) | 105 (Hartenfeller) |
| Action space building blocks (number & source) | 240,278 (Enamine REAL) |
| Action space building blocks embedding | Morgan fingerprint (1024) |

### C Training

#### C.1 Structure and confidence training

##### C.1.1 Main differences with Boltz-1

At a high level, the structure and confidence training phases are similar to those from Boltz-1, but there are a few key differences.

###### MSA sampling

In order to promote robustness of the model with regards to low-quality MSAs, at training time, we do not select MSA sequences greedily, but we rather sample them randomly among the top 16k hits. Moreover, given the recent progress on protein design models that rely on single-sequence folding predictions (without language model embeddings) [Pacesa et al., 2024, Cho et al., 2025], we aim at improving the model performance in this setting by randomly dropping all of the MSA of a complex in 5% of training iterations.

###### Ensemble supervision

While Boltz-1 supervises on only a single structure, Boltz-2 integrates multiple samples from experimental ensembles and MD trajectories. Structure supervision happens at two stages in the model: the trunk’s histogram output and the denoising module. Given an ensemble with *K* structures, we aggregate the one-hot encoded distograms of all *K* conformers and perform a weighted multi-class cross-entropy. For the coordinate noising and denoising supervision, we randomly sample at each training iteration one conformer to be used.

###### B-factor

On top of supervising the final pairwise representation of the trunk to predict the relative distances between pairs of tokens, we additionally supervise each token’s single representation to predict the B-factor of its representative atom. MD structures are supervised by computing the B-factor from the RMSF values computed over the trajectory, as given by [Kuzmanic et al., 2014]:

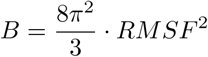

##### C.1.2 Controllability Training

Below is a description of the way that we sample templates, contact and pocket information at training time. Note that this sampling occurs after the cropping and therefore only applies to the entities and tokens that are present in the cropped structure.

###### Template sampling

Templates during training are sampled independently for each individual entity. 60% of the times no templates are selected for the selected chain. In the remaining 40% of the times, a random number of templates between 1 and min(1, # templates) is chosen. These templates are selected at random between the top 20 template hits for that particular entity. Multimeric templates are constructed by adding additional chains that are in the same PDB id and are template hits for chains in the complex that are still missing templates. No multimeric templating is done for symmetric entities to avoid issues with the chain mapping.

###### Contacts sampling

For each training example, we sample the number of contacts to add from a geometric distribution starting from 0 with *p* = 0.85. For each contact, we first sample the contact cutoff (maximum distance) between d=4^°^A and 20^°^A with a probability proportional to 1/d. Then, if there are multiple chains, we select a pair of tokens between different chains that have at least a pair of atoms within the cutoff. The sampling is done by selecting first a chain at random, enumerating all possible contacts and then selecting at random between the contacts. If there is only one chain, then we look for contacts between tokens that are at least 8 residues apart.

###### Pocket sampling

Similarly, to contact sampling, for each training example, we sample the number of pockets to specify from a geometric distribution starting from 0 with *p* = 0.85. For each pocket, we select a ligand at random if present, otherwise an arbitrary chain. Then, we sample the pocket cutoff (maximum distance) between d=4^°^A and 20^°^A with a probability proportional to 1/d. Then, the number of pocket tokens to specify for the pocket in question is sampled with a geometric distribution starting from 1 with *p* = 0.7. These pocket tokens are then sampled at random from the available ones.

##### C.1.3 Training stages

Table 5 shows the parameters used in the different stages of structure prediction training of the model. Most of the training happens at crop size of 384 tokens (atom crop size is always computed to be 9^∼^x the number of tokens) with more limited stages expanding this gradually to 512, 640, and 768. At the final stage, we exclude both the MD and the Boltz-1 distillation data to maintain only the highest quality datapoints. The molecular dynamics data was not included in the first stage of training due to project timing, we would expect bigger gains in the model’s ability to model dynamics had this data been integrated in the model earlier.

**Table 5:** Overview of the different stages for the structure prediction training.

| Training stage | Learning rate | Crop size | Training steps | Include MD | Include Boltz-1 distillation |
| --- | --- | --- | --- | --- | --- |
| 1st | 1e-3 | 384 | 88k | No | Yes |
| 2nd | 5e-4 | 512 | 4k | Yes | Yes |
| 3rd | 5e-4 | 640 | 4k | Yes | Yes |
| 4th | 5e-4 | 768 | 1k | No | No |

Confidence training was performed as a single stage using a crop size of 512 tokens and only trained on PDB data. In order to make the confidence model more robust to different inference hyperparameters, at every training iteration, we randomly sampled the number of inference steps between [20, 50, 200] and the diffusion step scale between [1.0, 1.1, 1.2, 1.3, 1.4, 1.5].

##### C.1.4 Architecture, Training and Inference Hyperparameters

Tables 6 and 7 record some of the hyperparameters of Boltz-2’s architecture, training and inference procedures that differ from Boltz-1’s and were not previously mentioned in the manuscript. For a full list of the hyperparameters and their precise impact on the model, we recommend the reader to refer directly to the code repository.

**Table 6:** Extra model architecture and training hyperparameters that differ from Boltz1 and were not previously mentioned in the manuscript.

| Parameter | Value |
| --- | --- |
| Max number of MSA sequences during training | 8192 |
| Template pairwise dim | 64 |
| Num template blocks | 2 |
| Training diffusion multiplicity | 32 |
| bfactor loss weight | $1 \times 10^{-3}$ |

**Table 7:** Diffusion process hyperparameters that differ from Boltz-1, with the exception of sigma min we opted for AlphaFold3’s default hyperparameters, see Abramson et al. [2024] for more details.

| Parameter | Value |
| --- | --- |
| sigma_min | 0.0001 |
| rho | 7 |
| gamma_0 | 0.8 |
| gamma_min | 1.0 |
| noise_scale | 1.003 |
| step_scale | 1.5 |

**Algorithm 2:**
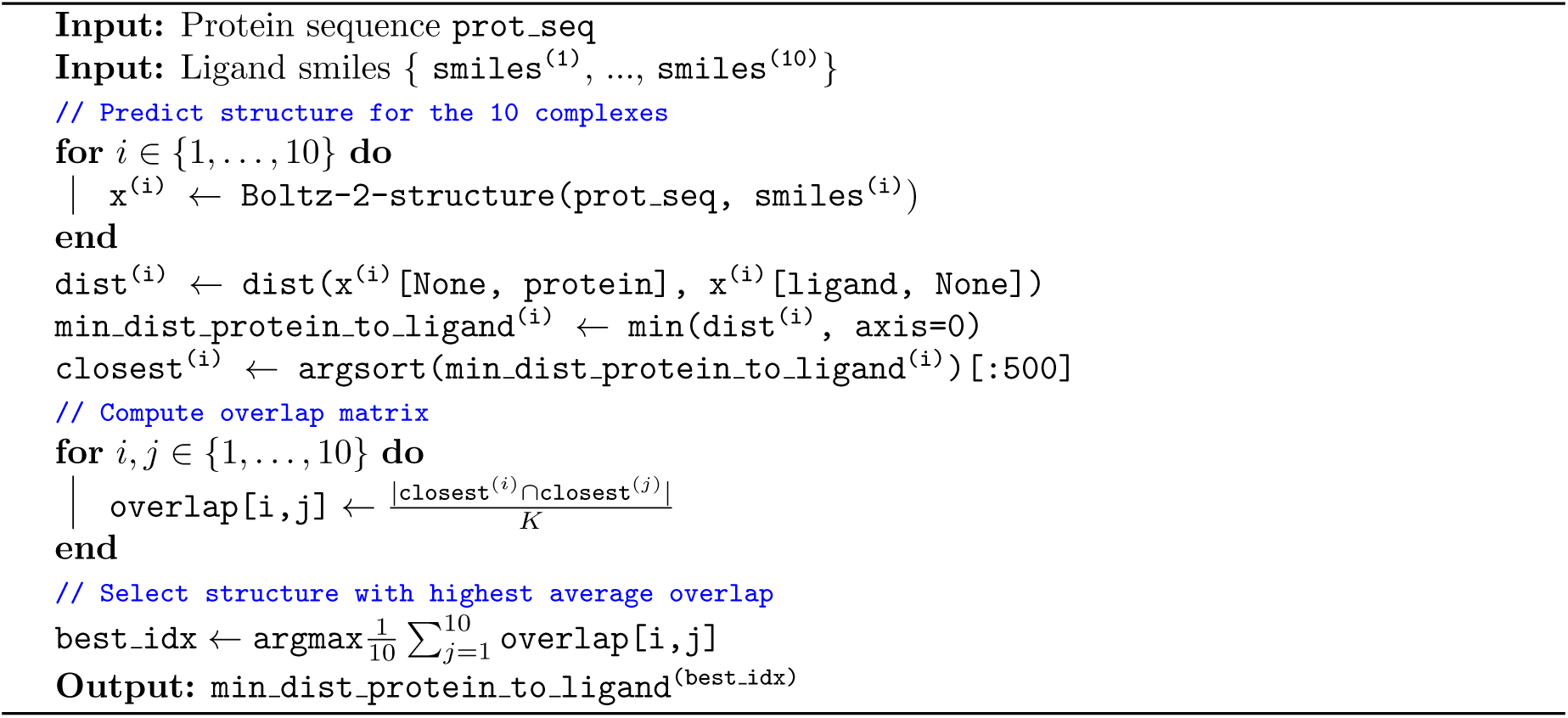
Pocket Pre-processing

#### C.2 Affinity training

Affinity training occurs after structural prediction, with gradients detached from the structural model to preserve its learned representations. Specifically, the training pipeline consists of the following key components:

1. Efficient pre-processing of protein binding pockets.
2. Cropping of spatial regions around the binding site.
3. Pre-processing of trunk features.
4. Sampling strategy that balances binders and decoys and prioritizes informative assays.
5. Robust loss functions tailored to mitigate the effects of experimental noise.

Each of these components is detailed in the following sections.

##### C.2.1 Pocket pre-processing

Target-based drug discovery generally assumes that most protein-ligand interactions occur within the binding pocket. To reduce the complexity of training and inference and reduce overfitting, we implement a pocket identification and cropping procedure.

The pocket is identified by computing the minimum atom-wise distance between the ligand and the surrounding protein structure. Specifically, for each target protein, we randomly sample 10 binders from the affinity training dataset and use Boltz-2 with 10 recycling steps, 200 diffusion iterations, and 5 structural samples per complex to predict their structure. We select the most confident structure according to the inter-chain predicted TM-score (ipTM).

From the resulting structures, we derive per-atom distance profiles between the protein and ligand. A consensus-based voting strategy is applied across the 10 binders’ structures to select the most likely binding site. The minimum distances from protein atoms to the ligand are cached and used by the *affinity cropper*. More details are provided in Algorithm 2.

##### C.2.2 Affinity cropper

We propose a cropping algorithm that leverages pre-computed pocket annotations to efficiently crop the complex around the binding site. This approach enables consistent cropping across complexes, even when all the complex structures are unavailable, and reduces the pre-processing complexity from O(# complexes) to O(# proteins).

**Algorithm 3:**
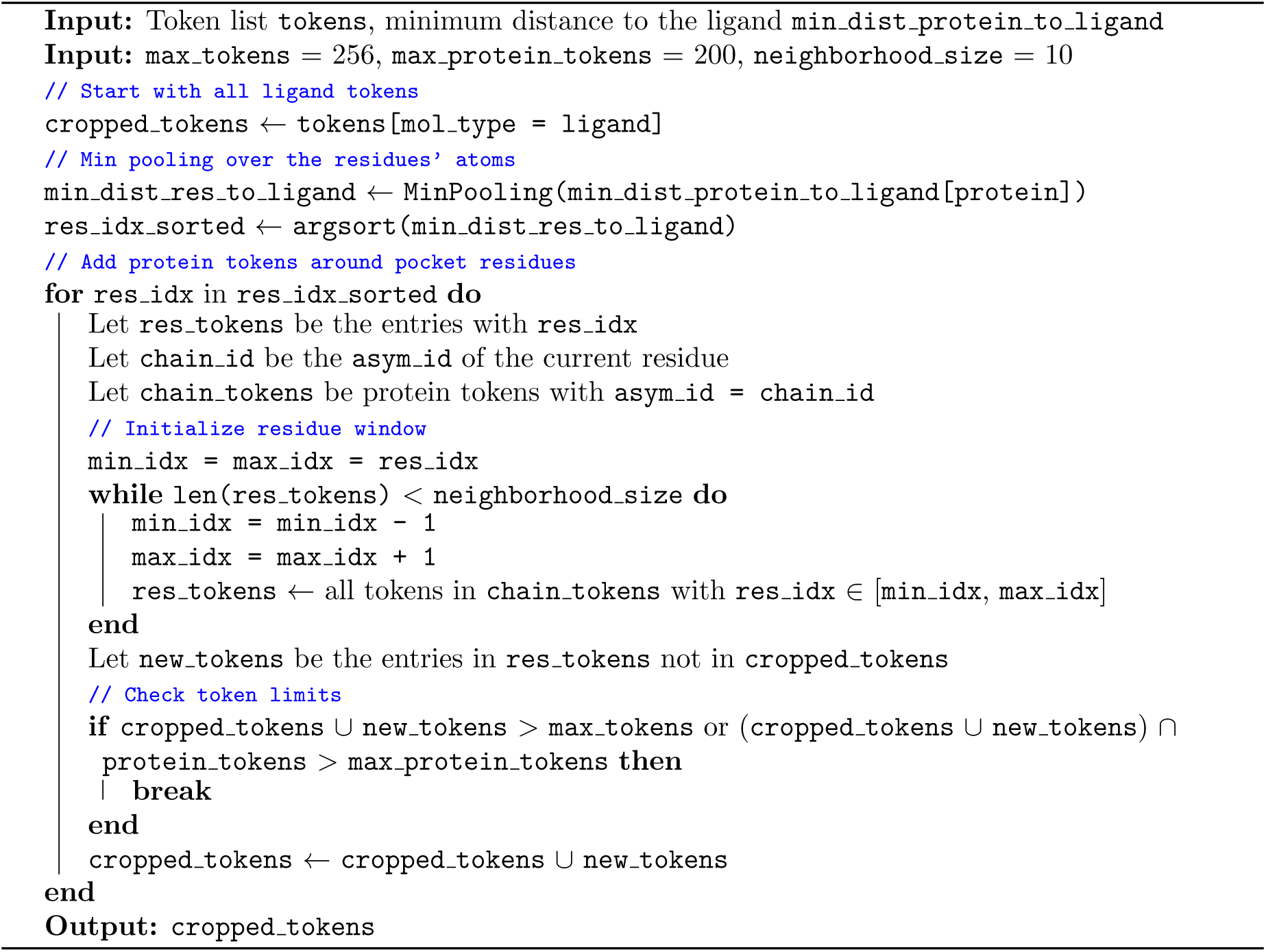
Affinity Cropper

The cropping procedure proceeds as follows. First, all ligand tokens are retained. Next, we apply a pocket-centered selection strategy inspired by the Boltz-2 structure model: For each protein token, we use the pocket annotation by selecting the nearest neighbors and apply the usual cropping algorithm with a neighborhood size of 10. We retain 256 tokens with a maximum of 200 protein tokens, to ensure consistency between molecules of different sizes at training.

Full details of the affinity cropper are provided in Algorithm 3.

##### C.2.3 Feature pre-processing

To reduce computational overhead during training, we pre-compute key structural and representational features. For each protein–ligand complex, we run Boltz-2 structure model with 5 recycling iterations, 200 diffusion steps, and generate 5 candidate structures. The most confident structure based on the interface predicted TM-score (ipTM) is retained for downstream use.

We extract and store the predicted atomic coordinates as well as the trunk pair representation and the cropped token indices. Since the affinity module utilizes only the protein–ligand and intra-ligand pairwise features, we discard the remaining pairwise interactions, and reduce the memory footprint by *>* 5x.

##### C.2.4 Training sampler

We design a custom affinity training sampler to enhance the model’s ability to learn from the noisy datasets. The sampler is constructed to balance binders and decoys, and to emphasize high-contrast assays that provide valuable learning signals.

**Algorithm 4:**
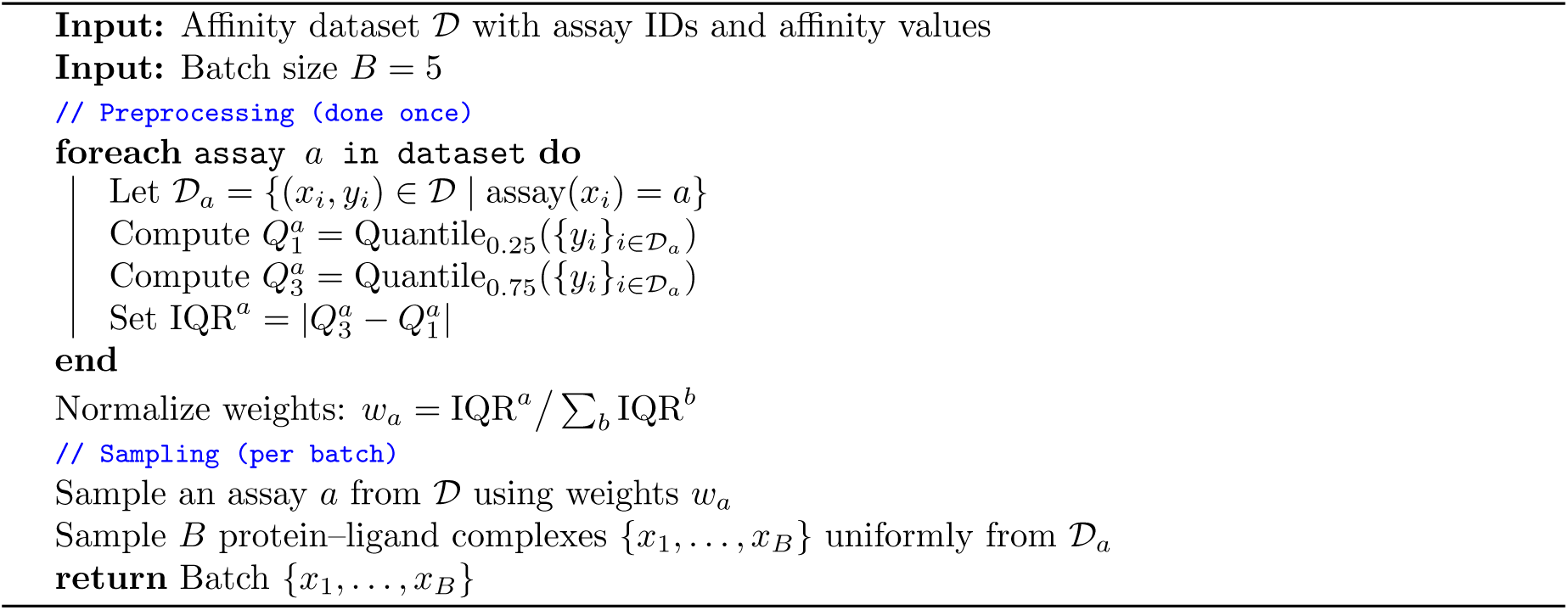
Activity Cliff Sampler

During training, we sample from the different data sources with probabilities specified in Table 8. For each sampled source, we construct batches of size 5, enforcing that all samples within a batch come from the same assay. We apply distinct sampling strategies depending on the type of label: datasets with continuous affinity values are treated differently from those with binary binding labels, allowing the model to better adapt to the nature of each supervision signal.

**Table 8:** Breakdown of affinity training data sources and their relative sampling weights.

| Source | Supervision | Sampling weight |
| --- | --- | --- |
| ChEMBL and BindingDB | values | 0.25 |
| PubChem small assays | values | 0.005 |
| PubChem HTS | binary | 0.44 |
| PubChem small assays | binary | 0.02 |
| CeMM Fragments | binary | 0.03 |
| MIDAS Metabolites | binary | 0.005 |
| ChEMBL and BindingDB synthetic decoys | binary | 0.25 |

###### Affinity value sampler

A key challenge in learning from affinity data lies in capturing activity cliffs—subtle, large shifts in binding affinity triggered by small structural modifications to a molecule. To encourage the model to focus on these high-frequency patterns, we sample five complexes coming from the same assay within each batch. This helps the model learn high-frequency variations and allows more complex loss functions as described in Appendix C.2.5.

To prioritize the most informative assays, we introduce an assay-level activity cliff score, defined as the interquartile range (IQR) of affinity values: the difference between the 75th and 25th percentiles of the affinity values. Sampling probabilities over assays are proportional to the activity cliff scores. Full details in Algorithm 4.

###### Binary label sampler

To improve discrimination between binders and decoys, we construct training batches with a consistent protein context. For each batch (1) we sample uniformly at random a binder from the dataset, (2) identify the associated assay and (3) randomly sample four decoys from the same assay.

##### C.2.5 Affinity supervision

We jointly train the binary classification and continuous affinity regression heads using the loss functions detailed below.

###### Affinity value supervision

Affinity measurements are notoriously noisy, with variability arising both from experimental replicates and inter-laboratory differences. Furthermore, *IC*_50_ values are highly sensitive to assay conditions, including substrate concentration and assay type, and may not be directly comparable across datasets [Landrum and Riniker, 2024]. While the Cheng–Prusoff equation is commonly used to convert *IC*_50_ to *K_i_*by correcting for substrate concentration, this correction is frequently infeasible due to missing metadata (eg. the substrate concentration or *K_m_*).

To address this, we introduce a supervision strategy based on pairwise differences of affinity values within the same assay. This difference-based formulation implicitly cancels out assay-specific confounding factors, such as those corrected by the Cheng–Prusoff equation.

We use a Huber loss—a quadratic loss for small errors *< δ*, but linear otherwise—to supervise both the affinity differences and the absolute values, with a weighted combination of the two. Many experimental affinity values are reported with inequality qualifiers (e.g., ‘=‘ or ‘*>*‘) rather than exact values. For entries with qualifier ‘*>*‘, we interpret the label as a lower bound and include the example in the loss only if the model prediction is *lower* than the reported value, both for the absolute affinity term and for its pairwise differences. This censor-aware supervision ensures that the model is not penalized for correct directional predictions when ground truth values are bounds rather than measurements. The resulting loss functions are:

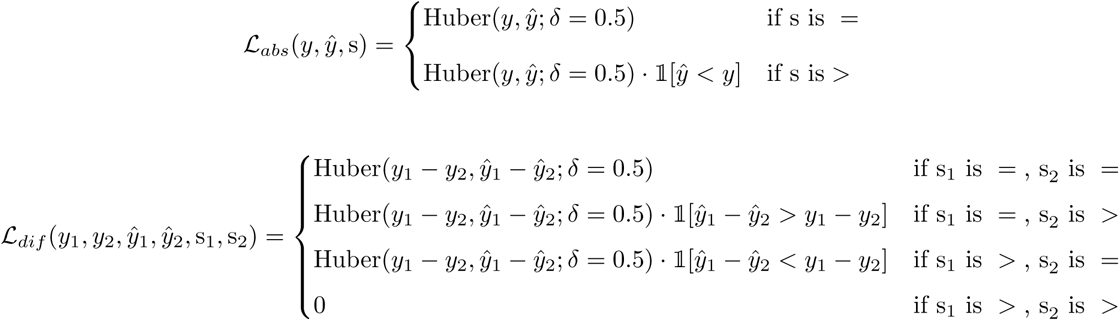

where s, s_1_ and s_2_ are the affinity qualifier of the ground truth measurement, and ]_ is the indicator function, which returns 1 if the condition holds, otherwise 0.

###### Binary label supervision

For binary binding classification, we use a focal loss with *γ* = 1, along with a balancing coefficient *λ_focal_* to weight the contribution of positive and negative samples:

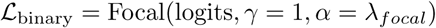

###### Overall loss

The final training objective is a weighted sum of the three components:

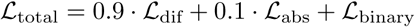

#### C.3 Generative ligand optimization with SynFlowNet

Binding affinity is among the most critical properties to optimize in the early stages of target-based drug discovery. Over the years, multiple generative models have been developed to generate novel ligands for early hit identification [Segler et al., 2018, Jensen, 2019, Du et al., 2024], but these methods often suffer from synthesizability issues and from the lack of a robust and fast scoring function to optimize against. Hence, generative models would often adversarially exploit the scoring function and generate non-sensical molecules, for example, by simply concatenating high-reward functional groups together [Renz et al., 2020, Langevin et al., 2022, Walters, 2024a,b].

Here we combine the proposed Boltz-2 model with SynFlowNet [Cretu et al., 2024], our recently published synthesis-aware molecular generator, to address these longstanding challenges. First, Boltz-2 offers the appropriate speed/accuracy trade-off needed for robust scoring functions. Our results in Section 5.3 show that Boltz-2 approaches FEP accuracy on public benchmarks. With an inference time of approximately 20 seconds per ligand, integrating Boltz-2 with modern parallel high-performance computing infrastructure enables the screening of hundreds of thousands of compounds per day. SynFLowNet, on the other end, offers the possibility of de novo molecular generation directly via synthetizable routes, naturally handling purchasability for downstream experimental validations. We combine these two models to perform a *Generative Virtual Screening* of ultra-large scale libraries of purchasable compounds (Enamine 76B REAL space) against a specific protein target. As opposed to Fixed-Library Virtual Screening, where a pre-assembled set of molecules is scored to identify potential hits, generative screens enable further exploration into the molecular space, far outside off-the-shelf libraries, by simultaneously training a parameterized sampler and building an annotated set of candidates (Figure 9).

The main score used for compound selection combines both the binding likelihood and the affinity value predictions of the first ensemble model (see Appendix B.5.2):

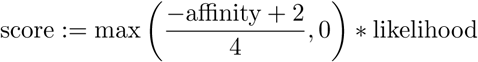

where the affinity value prediction is approximately normalised and lower-bounded to 0.

The SynFlowNet model uses this score as a reward function. The computational delay to obtain a numerical score for each molecule using Boltz-2 is approximately 20 seconds. With a batch size of 64, utilizing a single scoring instance would introduce significant latency in the training of the generative model. To mitigate this limitation, we implemented an asynchronous training paradigm: the SynFlowNet model continuously samples new trajectories from its current policy *P_F_* and submits them to a Last-In, First-Out (LIFO) Reward Queue. Concurrently, it samples batches of training data from an annotated Replay Buffer, which comprises all previously scored trajectories. An array of Boltz-2 scorers operates in parallel, retrieving new trajectories from the Reward Queue and depositing the annotated results into the Replay Buffer. A training batch pulled from the Replay Buffer consists of 50% trajectories uniformly sampled across the buffer (off-policy) and 50% of the newest trajectories added to the buffer (on-policy). For the screen presented in Section 5.4, we employed an array of 60 parallel Boltz-2 workers, each executing on a single H100 GPU. It is anticipated that the SynFlowNet policy will re-sample previously encountered molecules multiple times. To obviate unnecessary computations, we incorporated a caching system for the Boltz-2 workers, ensuring that scores are computed only for molecules newly added to the Replay Buffer. This caching system also serves to trigger the termination of the generative screen. Once a substantial majority of sampled molecules have been previously observed, indicating policy convergence, the generation of new trajectories is halted. In our specific case, this condition was met after approximately 400,000 molecules were sampled (corresponding to roughly 16 hours of runtime), totaling nearly 1,000 GPU-hours of computation for the generative screen.

### D Benchmarks and Baselines

#### D.1 Structure prediction

##### D.1.1 Benchmarks sets curation

We evaluate the models using recently released structures from the PDB, specifically from 01/01/24 to 12/31/2024. We filter the structures such that a target is kept if it has at least one monomer that is more than 40% sequence dissimilar to chains in the training data. For interfaces, we keep only interfaces where at least one of the two chains is sufficiently distinct, as defined above. In addition, we remove targets with resolution worse than 4.5A and with more than 1200 residues. This yields a final set of 2315 unique targets, from which we evaluate the subset of novel chains and interfaces.

##### D.1.2 Evaluation metrics

We evaluate the models using two sets of metrics:

- The lDDT (local Distance Difference Test), which we compute per modality (protein, RNA, DAN, ligands) as well as on specific interfaces of any two modalities.
- DockQ which we use to score antibody interfaces.

For each result, we provide the top-1 prediction across 5 samples according to the confidence model ranking.

For MD evaluation, we report the following metrics:

1. lDDT (local Distance Difference Test)

a. Precision: the average lDDT score from each predicted structure to its closest corresponding crystal structure.
b. Recall: the average lDDT score from each crystal structure to its closest predicted structure.
c. Diversity: the average structural dissimilarity between pairs of predicted structures, calculated as 1 − *lDDT* .
2. RMSF-*Cα* (root mean square fluctuation): measuring the atom-level flexibility over all conformations of the MD trajectory, and measuring its correlation to the generated data:

Spearman’s rank correlation coefficient (*ρ*), pooled globally (by first aggregating all RMSF values across targets and then computing correlation metrics) as well as locally (by first computing the correlation within each target and then taking the median value across the dataset).
Pearson’s correlation coefficient (*r*), also pooled globally and locally.
Root Mean Squared Error (RMSE) over the predicted and reference RMSF values, both locally and globally.

MD metrics were computed by taking 100 samples from Boltz-2 and other baseline models against 200 frames from the ATLAS and mdCATH trajectories.

##### D.1.3 Baselines

###### PDB baselines

We evaluate Boltz-2 structural prediction results on the PDB against other state-of-the-art co-folding models, AlphaFold3, Chai-1, ProteinX, and Boltz-1. For all tools, we use the same inference parameters (5 recycling rounds, 5 samples, single seed) and the same MSA. We do not use templates during evaluation.

###### MD baselines

To evaluate the ability of Boltz-2 to model multiple conformations, we compare against Boltz-1 [Wohlwend et al., 2025], AlphaFlow-MD base [Jing et al., 2024] and BioEmu with HPacker for side-chain reconstruction without MD relaxation [Lewis et al., 2025]. AlphaFlow is excluded from the ATLAS evaluation given that its training set largely overlaps with the test set constructed for Boltz-2. All models were run with the same MSA and sequence inputs.

#### D.2 Affinity prediction

This section will detail all the steps used to evaluate the affinity prediction. The evaluation addresses both hit discovery binary prediction, as well as the ranking of chemical series for hit-to-lead and lead optimization stages.

##### D.2.1 Benchmark sets curation

To rigorously evaluate the performance of our model across diverse binding tasks, we construct a curated suite of benchmark datasets targeting both continuous affinity value prediction and binary classification. These benchmarks are designed to reflect real-world drug discovery challenges, including hit-to-lead optimization and high-throughput screening.

###### Affinity value benchmarks

For the affinity regression task, we curate a validation set by selecting a diverse collection of hit-to-lead assays from our training corpus. The assays are chosen according to the following criteria:

- We retain only *K_i_* measurements to ensure a higher degree of experimental consistency and lower measurement noise.
- Assays must exhibit a sufficiently large dynamic range in affinity values, enabling the model to distinguish strong from weak binders.
- We exclude assays with high correlation between affinity and molecular weight to minimize artifacts introduced by molecular series subselection.
- The selected assays span a diverse set of protein families to ensure broad generalization.

This filtering process yields 16 assays drawn from BindingDB and ChEMBL:

- ChEMBL assay ids: 1528727, 438257, 1572912, 1705740, 157530, 454476, 2114176, 1527798, 769558.
- BindinDB DOIs: 10.7270/Q2JQ0ZNX, 10.7270/Q2ZC81K5, 10.7270/Q2VX0FFW, 10.7270/Q2RV0MMK, 10.7270/Q2VD71JR, 10.7270/Q26D5WBR, 10.7270/Q2VD72PZ.

We evaluate model performance on the following held-out affinity test sets:

- 2 subsets of the FEP+ benchmark: The OpenFE subset consisting of 876 protein–ligand complexes from hit-to-lead and lead optimization campaigns as well as a 4 targets subset (CDK2, TYK2, JNK1, P38; curated in the protein-ligand-benchmark [Hahn et al., 2022]) with 87 neutral compounds [Chen et al., 2023].
- A proprietary collection of internal hit-to-lead assays provided by Recursion.
- The CASP16 binding affinity challenge dataset.

###### Binary classification benchmarks

For binary binding prediction, we construct a validation set using six biochemical assays from the MF-PCBA dataset.

For the final test set, we select 10 biochemical high-throughput screening (HTS) assays from the MF-PCBA benchmark to maximize functional diversity. The following filtering steps are applied:

- We parse data directly from the MF-PCBA GitHub repository, adopting their assay-specific subselection strategy (e.g., retaining only ligands labeled as active in secondary confirmatory assays).
- Each assay is randomly downsampled to 50,000 protein–ligand complexes.
- We remove all compounds flagged as PAINS (Pan-Assay Interference Compounds) to mitigate false-positive artifacts.

The selected PubChem assay identifiers used for the test sets are: 743445, 485317, 2097, 493091, 2650, 485273, 504329, 489030, 588689, 588549.

###### Data leakage control

To avoid information leakage between training and evaluation splits, we apply strict sequence-level filtering. Specifically, we exclude from the training set any proteins with sequence similarity ≥ 90% to proteins in the validation or test sets. This is implemented by first clustering all protein sequences in the affinity datasets using ‘mmseqs easy-cluster … –min-seq-id 0.9 – cov-mode 0 -c 0.01‘ [Hauser et al., 2016], and then removing any training protein that falls into a cluster shared with a validation or test protein. This filtering is applied to all benchmark datasets except for CASP16 and the Recursion internal assays: CASP16 data was released after our training data cutoff, and the Recursion benchmarks consist of proprietary internal targets not accessible to external sources. Moreover, for the FEP+ benchmark, we assess the impact of compound similarity in Figure D.2.1 by computing the maximum Tanimoto similarity of each test compound to the affinity value training set, followed by mean pooling across assays. We observe no significant dependence between prediction performance and compound similarity. We perform the same analysis on the CASP16 benchmark, obtaining maximum Tanimoto similarities of 0.41 for the L1000’s compounds and 0.59 for the L3000’s compounds, both sufficiently low to alleviate concerns about compound-level information leakage.

###### Benchmarking challenges

Improving benchmark design is essential for advancing affinity prediction across hit discovery, hit-to-lead, and lead optimization applications. Many widely used benchmarks employ subselection strategies that introduce artificial biases and obscure the challenges inherent in real-world campaigns. In binary activity datasets, noise from off-target effects and surrogate readouts further complicates learning and evaluation, often enabling models to exploit dataset artifacts rather than true structure–activity relationships. To assess model performance more meaningfully, we advocate for the creation of standardized benchmarks and careful curation to reduce experimental noise and spurious correlations, offering more reliable test sets for models intended to support early-stage compound prioritization and lead refinement.

##### D.2.2 Evaluation metrics

We evaluate our model on both continuous affinity prediction and binary binder classification tasks.

###### Affinity Value Prediction

All regression metrics are calculated per assay and averaged across assays with weight proportional to the number of compounds in each assay, ensuring that larger assays contribute proportionally to the overall performance summary. All predictions are converted to kcal/mol prior to computing the metrics. For the regression task, we report the following metrics:

1. Pearson’s correlation coefficient (R), to measure linear correlation between predicted and true binding affinities.
2. Kendall’s Tau (*τ*) rank correlation coefficient, to assess the monotonic agreement between predicted and experimental affinities within each assay.
3. Pairwise Mean Absolute Error (PMAE), calculated as the MAE over the pair-wise difference of affinity between any pair of compounds in a given assay.
4. Mean Absolute Error (MAE), computed between predicted and measured affinity values.
5. Percentage within 1 and 2 kcal/mol, indicating the fraction of predictions with absolute error below 1 and 2 kcal/mol, respectively.

For PMAE, MAE, and PW, we additionally report a *centered* version in which the predicted affinity values for each assay are translated to have the same mean as the corresponding ground truth values.

This adjustment allows for a fair comparison with methods that only predict relative affinities, such as relative FEP.

###### Binary label Prediction

For the binary classification task, we compute the metrics per assay and average them uniformly across assays regardless of size. We report the following metrics:

1. Average Precision (AP), corresponding to the area under the Precision-Recall curve.
2. Enrichment Factor (EF) at top 0.5 %, 1%, 2% and 5% of the ranked list of compounds, reflecting early retrieval performance.
3. Area Under the Receiver Operating Characteristic Curve (AUROC), evaluating the model’s ability to distinguish binders from non-binders.
4. *global* AUROC, where the metric is computed across the full set of compounds and targets jointly, rather than per-assay.

##### D.2.3 Baselines

Existing work for binding affinity prediction can be categorized into physics-based or ML-based approaches. Physics-based approaches include a range of methods across the cost-accuracy Pareto front, ranging from inexpensive Docking and QM-based scoring functions, through MD-based endstate approaches such as MM/PBSA to physically rigorous (alchemical) free energy simulations such as absolute and relative FEP. ML-based approaches can be roughly classified into structure-based and sequence-based methods. The former require the 3D (crystal) structure of the protein-ligand complex and are therefore usually only trained on the 20k complexes in PDBBind [Wang, 2024, Zhang et al., 2023]. The latter only require knowledge about the compound SMILES and protein sequence and can therefore be trained on millions of experimental binding measurements without associated structures.

As an ML baseline, we select the sequence-based method *BACPI* [Li et al., 2022], given that the vast majority of our training data does not include structural information. BACPI consists of a 1D convolutional NN that processes the protein sequence and a graph attention transformer (GAT) that processes the ligand SMILES, connected by bi-directional attention. This baseline allows us to estimate the performance gain from leveraging predicted structural information compared to sequence-information only. In addition, we train ligand-only models to estimate the ligand bias in the data by deactivating the protein sequence CNN of BACPI (referred to as GAT in the following). We select the default hyperparameters of the original publication [Li et al., 2022], except for the batch size, which we increased to 32 to speed-up training on significantly larger datasets.

We also compare the performance of Boltz-2 to the following physical baselines: For ranking congeneric compounds in a hit-to-lead setting, we use our recently-published ABFE protocol [Wu et al., 2025] as well as the relative FEP protocols of OpenFE [Gowers et al., 2023] and FEP+ [Wang et al., 2015] for benchmarking. The results of FEP+ are intended to show the maximum attainable accuracy of current (commercial) FEP simulations by manually adjusting the protocol (input preparation, perturbation map, force field) to the system at hand after observing the error with respect to the experiment [Ross et al., 2023]. In addition, the results of the ABFE protocol [Wu et al., 2025] on the 4 target subset may also represent an optimistic estimate of its prospective performance given that the protocol was optimized based this dataset. In contrast, the OpenFE results are representative of automated open-source simulations (fixed protocol) [Horton, 2025]. We also compare to less expensive physics-based scoring functions based on docked poses obtained by Glide [Friesner et al., 2004] in the FEP+ dataset [Ross et al., 2023]. These include OpenEye’s Chemgauss4 and an in-house DFTB3-based Fragment Molecular Orbital (FMO) code [Nishimoto and Fedorov, 2016, Guareschi et al., 2023]. As a representative of endstate approaches, we select the MM/PBSA implementation of AMBER [Miller III et al., 2012, Case et al., 2023]: Initial modeled complexes were thermalized with a position-restrained minimization and equilibration procedure, finishing with a 1 ns unrestrained simulation under NPT conditions. The final frame from this simulation was used for MM/PBSA scoring. In the binary hit discovery setting, we select the commercial Docking engine OpenEye FRED [McGann, 2011] to predict binding poses. Any undefined R/S or E/Z stereocenters in input molecules were exhaustively enumerated and molecular conformers for FRED docking were generated with OpenEye’s OMEGA with default sampling options. We rank compounds by highest Chemgauss4 efficiency: Chemgauss4 score squared, divided by the number of heavy atoms. In the case of multiple stereoisomers per input molecule, we select the isomer with the highest score. In the absense of experimental crystal structures in MF-PCBA, we co-fold the median-weight active compound of each assay with Boltz-2 to obtain the receptor structure for Docking. In all cases, we prepare protein structures with OpenEye Spruce.

### E Extended Results

In this section, we present additional results, evaluation metrics, and analyses covering the various components of the model. Due to the scale of Boltz-2, comprehensive ablation studies isolating the impact of each architectural or training component on final performance are not computationally feasible.

Note that this paper is still a preprint in preparation. In the coming weeks, we will integrate even more results to the paper including evaluations of the template, contact and pocket conditioning as well as more challenging prospective evaluations of our small-molecule design pipelines.

#### E.1 Molecular Dynamics

Tables 9 and 10 present the precise measurements of the RMSF correlations and RMSE of the various models on a per-target and global basis for holdout MD datasets. Figure 11 displayes the global correlations between predicted and groundtruth RMSF values.

**Table 9:** mdCATH test set. Comparison of methods based on RMSF metrics: correlation (*r*), Spearman’s rank correlation (*ρ*), and mean squared error (MSE), both globally and per target. Boltz-2 is run with MD and X-ray method conditioning.

| Metric | Boltz-2 - Xray | Boltz-2 - MD | Boltz-1 | AlphaFlow | BioEmu |
| --- | --- | --- | --- | --- | --- |
| ↑ Global RMSF $r$ | 0.48 | **0.67** | 0.46 | 0.24 | 0.53 |
| ↑ Per-target RMSF $r$ | 0.72 | **0.79** | 0.70 | 0.77 | 0.77 |
| ↑ Global RMSF $\rho$ | 0.61 | **0.65** | 0.52 | 0.45 | 0.44 |
| ↑ Per-target RMSF $\rho$ | 0.78 | **0.81** | 0.76 | 0.76 | 0.78 |
| ↓ Global RMSF RMSE | 192 | **157** | 197 | 229 | 212 |
| ↓ Per-target RMSF RMSE | 21.71 | 16.30 | 22.92 | 18.74 | **14.85** |

**Table 10:** ATLAS test set. Comparison of methods based on RMSF metrics: correlation (*r*), Spearman’s rank correlation (*ρ*), and mean squared error (MSE), both globally and per target. Boltz-2 is run with MD and X-ray method conditioning.

| Metric | Boltz-2 - Xray | Boltz-2 - MD | Boltz-1 | BioEmu |
| --- | --- | --- | --- | --- |
| ↑ Global RMSF $r$ | 0.57 | **0.65** | 0.38 | 0.56 |
| ↑ Per-target RMSF $r$ | 0.76 | **0.85** | 0.77 | 0.83 |
| ↑ Global RMSF $\rho$ | 0.63 | **0.76** | 0.67 | 0.63 |
| ↑ Per-target RMSF $\rho$ | 0.82 | **0.87** | 0.83 | 0.81 |
| ↓ Global RMSF RMSE | 185 | **155** | 218 | 209 |
| ↓ Per-target RMSF RMSE | 17.42 | **12.35** | 19.62 | 15.04 |

#### E.2 Affinity prediction

##### E.2.1 Public benchmarks

In this section, we provide the complete set of quantitative results for the model’s performance across all public benchmarks used in our evaluation. These are intended to supplement the main text by offering a more detailed view of the metrics and trends observed in each dataset.

Tables 11 and 12 present the comprehensive benchmark results for the FEP+ dataset, including both OpenFE subset and the focused 4-target evaluation. Table 15 shows the model’s performance on our affinity values validation set, which was constructed to span a representative set of high-quality hit-to-lead assays. In Figure 12, we visualize the predicted versus measured affinity values for each assay in the validation set. Each scatter plot is annotated by protein class (e.g., kinases, GPCRs, proteases), enabling visual assessment of potential systematic trends. While we do not observe any systematic degradation or improvement in performance attributable to specific protein classes, we do find substantial variability across assays, with Pearson correlations ranging from 0.732 to 0.056. Table 14 reports Boltz-2’s results on the CASP16 blind challenge, alongside the ML baselines and the top-6 ranked entries from the competition. Lastly, Table 13 presents the complete evaluation metrics for the MF-PCBA test set, while Figure 13 complements this by showing ROC curves and activity histograms for individual assays.

**Table 11:** OpenFE subset of the FEP+ benchmark. Comparison of Boltz-2, ML baselines and free energy perturbation.

| method | type | Pearson R<br>target avg. | Kendall tau<br>target avg. | PMAE<br>target avg. | MAE |  | Perc. within 1 |  | Perc. within 2 |  |
| --- | --- | --- | --- | --- | --- | --- | --- | --- | --- | --- |
|  |  |  |  |  | non-cent. | cent. | non-cent. | cent. | non-cent. | cent. |
| **Boltz-2** | ML | 0.62 | 0.46 | 0.93 | 1.22 | 0.64 | 0.49 | 0.80 | 0.82 | 0.96 |
| Boltz-2 iptm | ML | -0.07 | -0.05 | N/A | N/A | N/A | N/A | N/A | N/A | N/A |
| GAT | ML | 0.28 | 0.20 | 1.30 | 1.42 | 0.91 | 0.40 | 0.64 | 0.75 | 0.92 |
| BACPI | ML | 0.29 | 0.19 | 1.21 | 1.44 | 0.85 | 0.40 | 0.67 | 0.74 | 0.94 |
| OpenFE | physics | 0.63 | 0.47 | 1.37 | N/A | 0.94 | N/A | 0.65 | N/A | 0.91 |
| FEP+ | physics | 0.72 | 0.53 | 0.94 | N/A | 0.64 | N/A | 0.79 | N/A | 0.97 |

**Table 12:** 4 target subset of the FEP+ benchmark. Comparison of Boltz-2, ML baselines and an extensive set of physics-based methods.

| method | type | time | Pearson R<br>target avg. | Kendall tau<br>target avg. | PMAE<br>target avg. | MAE |  | Perc. within 1 |  | Perc. within 2 |  |
| --- | --- | --- | --- | --- | --- | --- | --- | --- | --- | --- | --- |
|  |  |  |  |  |  | non-cent. | cent. | non-cent. | cent. | non-cent. | cent. |
| **Boltz-2** | ML | 20 GPU sec | 0.66 | 0.48 | 0.85 | 0.75 | 0.59 | 0.69 | 0.83 | 0.97 | 0.98 |
| Boltz-2 iptm | ML | 5 GPU sec | 0.04 | 0.09 | N/A | N/A | N/A | N/A | N/A | N/A | N/A |
| BACPI | ML | 0.48 GPU ms | 0.14 | 0.09 | 1.18 | 1.40 | 0.82 | 0.43 | 0.62 | 0.73 | 1.00 |
| GAT | ML | 0.18 GPU ms | 0.40 | 0.28 | 1.07 | 1.19 | 0.71 | 0.43 | 0.72 | 0.86 | 0.95 |
| OpenFE | physics | 6-12 GPU hours | 0.66 | 0.51 | 1.09 | N/A | 0.75 | N/A | 0.70 | N/A | 0.98 |
| FEP+ | physics | - | 0.78 | 0.63 | 0.77 | N/A | 0.53 | N/A | 0.85 | N/A | 1.00 |
| ABFE | physics | >20 GPU hours | 0.75 | 0.54 | 0.95 | 2.47 | 0.65 | 0.11 | 0.79 | 0.40 | 0.98 |
| FMO | physics | 2-10 CPU min | 0.55 | 0.38 | N/A | N/A | N/A | N/A | N/A | N/A | N/A |
| MM/PBSA | physics | 10-15 CPU min | 0.18 | 0.16 | N/A | N/A | N/A | N/A | N/A | N/A | N/A |
| Chemgauss4 | physics | 20-30 CPU sec | 0.26 | 0.17 | N/A | N/A | N/A | N/A | N/A | N/A | N/A |

**Table 13:** MF-PCBA test set. Comparison of Boltz-2 with ML baselines, confidence score and Chemgauss4 Docking score.

| method | AP<br>target avg. | EF at 0.5%<br>target avg. | EF at 1%<br>target avg. | EF at 2%<br>target avg. | EF at 5%<br>target avg. | AUROC |  |
| --- | --- | --- | --- | --- | --- | --- | --- |
|  |  |  |  |  |  | target avg. | global |
| **Boltz-2** | 0.0248 | 18.3916 | 13.9540 | 10.5706 | 7.0448 | 0.8122 | 0.8056 |
| Chemgauss4 | 0.0051 | 1.9969 | 2.2257 | 2.1136 | 1.6462 | 0.5450 | 0.5706 |
| Boltz-2 iptm | 0.0046 | 2.4242 | 3.1728 | 2.6881 | 2.2263 | 0.5657 | 0.6134 |
| GAT | 0.0133 | 11.1179 | 8.9897 | 7.5630 | 5.9055 | 0.7928 | 0.7867 |
| BACPI | 0.0131 | 9.4818 | 9.2397 | 7.3983 | 5.5533 | 0.7575 | 0.7205 |

**Table 14:** CASP16 competition. Comparison of Boltz-2 and the ML baselines with the top-6 highest ranked participants.

| method | Pearson R | Kendall tau | Pairwise MAE | MAE |  | Perc. within 1 |  | Perc. within 2 |  |
| --- | --- | --- | --- | --- | --- | --- | --- | --- | --- |
|  | target avg. | target avg. | target avg. | non-cent. | cent. | non-cent. | cent. | non-cent. | cent. |
| **Boltz-2** | 0.65 | 0.45 | 1.36 | 1.28 | 0.95 | 0.48 | 0.61 | 0.81 | 0.90 |
| LG016 | 0.54 | 0.42 | 1.43 | 1.09 | 1.03 | 0.53 | 0.51 | 0.83 | 0.91 |
| LG082 | 0.38 | 0.36 | 1.55 | 1.17 | 1.12 | 0.49 | 0.47 | 0.83 | 0.85 |
| LG204 | - | 0.34 | - | - | - | - | - | - | - |
| LG055 | 0.47 | 0.33 | 1.59 | 1.29 | 1.12 | 0.45 | 0.53 | 0.78 | 0.84 |
| LG207 | 0.38 | 0.32 | 1.65 | 1.32 | 1.19 | 0.48 | 0.47 | 0.74 | 0.83 |
| LG008 | 0.38 | 0.29 | 1.62 | 1.43 | 1.16 | 0.39 | 0.51 | 0.72 | 0.85 |
| Boltz-2 iptm | 0.12 | 0.07 | N/A | N/A | N/A | N/A | N/A | N/A | N/A |
| GAT | 0.50 | 0.35 | 1.58 | 1.28 | 1.13 | 0.44 | 0.49 | 0.79 | 0.84 |
| BACPI | 0.41 | 0.31 | 1.55 | 1.25 | 1.10 | 0.45 | 0.51 | 0.81 | 0.89 |

**Table 15:** Hit-to-lead affinity validation set. Comparison between Boltz-2 and ML baselines.

| method | Pearson R | Kendall tau | PMAE | MAE |  | Perc. within 1 |  | Perc. within 2 |  |
| --- | --- | --- | --- | --- | --- | --- | --- | --- | --- |
|  | target avg. | target avg. | target avg. | non-cent. | cent. | non-cent. | cent. | non-cent. | cent. |
| **Boltz-2** | 0.4246 | 0.2855 | 1.2046 | 1.7001 | 0.8569 | 0.3071 | 0.6555 | 0.6223 | 0.9351 |
| GAT | 0.2512 | 0.1795 | 1.3117 | 1.4261 | 0.9308 | 0.4288 | 0.6231 | 0.7433 | 0.9043 |
| BACPI | 0.1997 | 0.1330 | 1.2855 | 1.8775 | 0.9103 | 0.2913 | 0.6260 | 0.5861 | 0.9120 |

**Table 16:** Benchmark on 8 blinded, private hit-to-lead binding assays. Comparison of Boltz-2 with ML baselines.

| method | Pearson R | Kendall tau | Pairwise MAE | MAE |  | Perc. within 1 |  | Perc. within 2 |  |
| --- | --- | --- | --- | --- | --- | --- | --- | --- | --- |
|  | target avg. | target avg. | target avg. | non-cent. | cent. | non-cent. | cent. | non-cent. | cent. |
| **Boltz-2** | 0.39 | 0.23 | 1.91 | 1.67 | 1.36 | 0.38 | 0.45 | 0.66 | 0.77 |
| GAT | 0.16 | 0.11 | 2.16 | 1.84 | 1.47 | 0.32 | 0.41 | 0.61 | 0.73 |
| BACPI | 0.11 | 0.06 | 2.11 | 1.87 | 1.44 | 0.32 | 0.42 | 0.59 | 0.74 |

**Table 17:** Number of compounds after each filtering stage of the screening pipeline for all five streams.

| Stream | Initial | Score > 0.5 | in REAL | MPO 1&2 | Max Diversity | Final |
| --- | --- | --- | --- | --- | --- | --- |
| SynflowNet screen | 117,199 | → 16,317 | → 1,996 | → 93 | → 10 | → 10 |
| Enamine HLL screen | 460,160 | → 239 | → - | → 10 | → - | → 10 |
| Enamine Kinase Lib screen | 64,960 | → 506 | → - | → 10 | → - | → 10 |
| Random REAL sample | 1,000 | → - | → 1,000 | → - | → 10 | → 10 |
| Public TYK2 binders | 10 | → - | → - | → - | → - | → 10 |

##### E.2.2 Private benchmarks

This section supplements the main text discussion about the benchmarking of Boltz-2 on blinded, past experimental binding assays performed at Recursion, which represents a more rigorous evaluation of the types of problems we expect Boltz-2 to be exposed to in real-world drug discovery projects. Each assay has hundreds of compounds that were screened during the hit-to-lead stage. Unlike the deep learning baselines, Boltz-2 still achieves decent correlation with the experiment on average (table E.2.2). As displayed in figure 14, the model achieves respectable performance on these assays, achieving an average Pearson R = 0.39, only slightly worse than in the validation set (R = 0.42). However, the centered MAE = 1.36kcal/mol is significantly worse compared to the validation set (MAE = 0.86kcal/mol). In addition, the performance varies noticably between targets, ranging from Person R = 0.165 to R = 0.634 and centered MAE from MAE = 0.855kcal/mol to MAE = 1.734kcal/mol, suggesting that performance in practice will strongly depend on the project at hand. These results highlight challenges in real-world drug discovery projects that may be insufficiently reflected in public benchmarks. This effect has been recently observed for benchmarking OpenFE as well, where performance significantly dropped on private data compared to the FEP+ benchmark [Horton, 2025].

#### E.3 Prospective virtual screens

In this section, we provide addtionnal details and results from the filtering pipeline of the prospective virtual screens of Section 5.4, visualisations of all of the ABFE-screened molecules from Figure 8, 3D renderings of the co-folded structure of the top two ligands (as ranked by ABFE) with the TYK2 protein, and a similarity analysis with the TYK2 ligands that were contained in Boltz-2 structural training data through the PDB.

##### E.3.1 Ligand filtering pipeline

Table E.3.1 shows the number of remaining compounds after each filtering stage of our screening pipeline, for both our generative SynFlowNet screen and the fixed virtual screens of Enamine’s HLL and Kinase libraries. The first and most important stage is computing the scores using Boltz-2 for each of the compounds under consideration. For SynFlowNet, 117, 199 unique compounds were scored out of the 400*k* samples from the model throughout the training of the model. We sequentially (1) removed all compounds with a score below 0.5, (2) discarded all the compounds not contained in the REAL space, to guarantee purchasability, (3) performed multi-parameter optimization (MPO) the scores computed by both ensemble models to be above 0.9, and (4) enumerated all undefined R/S or E/Z stereocenters of the selected molecules and removed compounds with 4 or more stereoisomers to reduce the ABFE simulation cost. To finalise the candidate set for ABFE-validation, we selected 10 diverse compounds among the 93 remaining candidates using Tanimoto fingerprint similarity. For fixed libraries (HLL and Kinase), we notice that since these libraries were not assembled specifically to optimise against our target, contrary to the SynFlowNet stream, we are left with only a few hundreds remaining compounds after imposing a score threshold of 0.5. The compounds being already purchasable, we do not test containment to the REAL space and simply select the set of 10 compounds that provide the best joint scores across the ensemble. At this point, not enough samples were left to further maximise diversity. The results in Figure 15 show how the score distributions of the selected compounds differ from a random control set and a set of public TYK2 binders from the protein-ligand benchmark Hahn et al. [2022].

**Figure 15:**
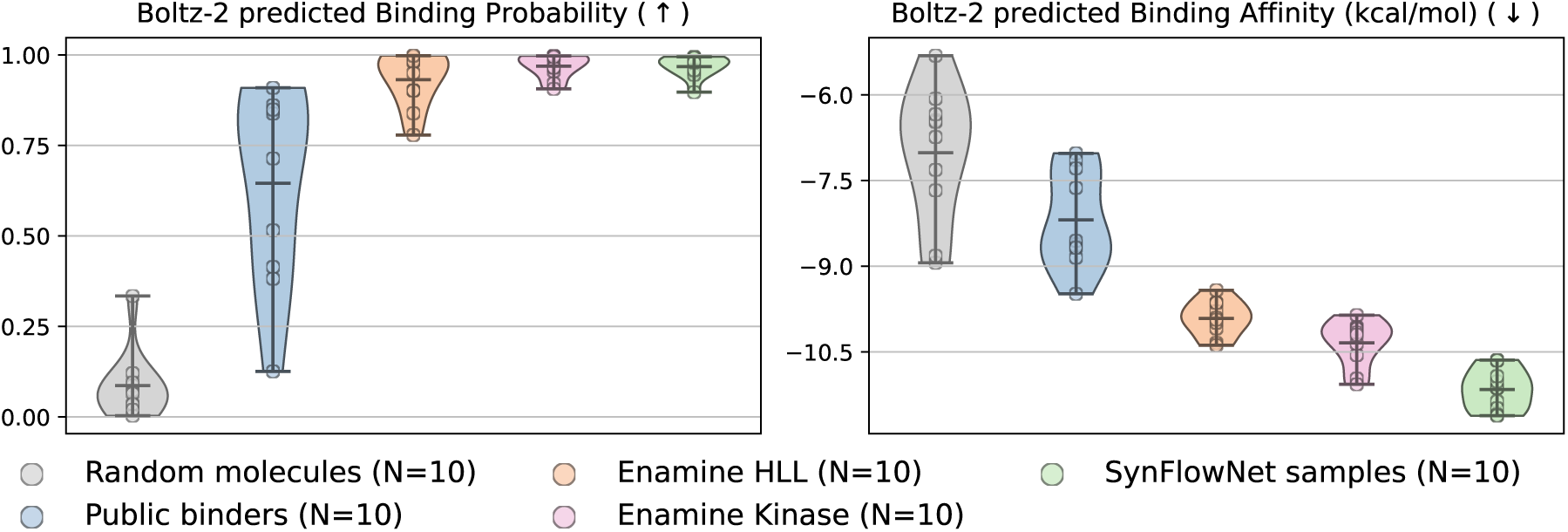
In virtual screening, Boltz-2 identifies what it believes to be high-scoring compounds, with the SynFlowNet obtaining higher scores on average. These potential binders are then tested with ABFE (results in Figure 8).

**Figure 16:**
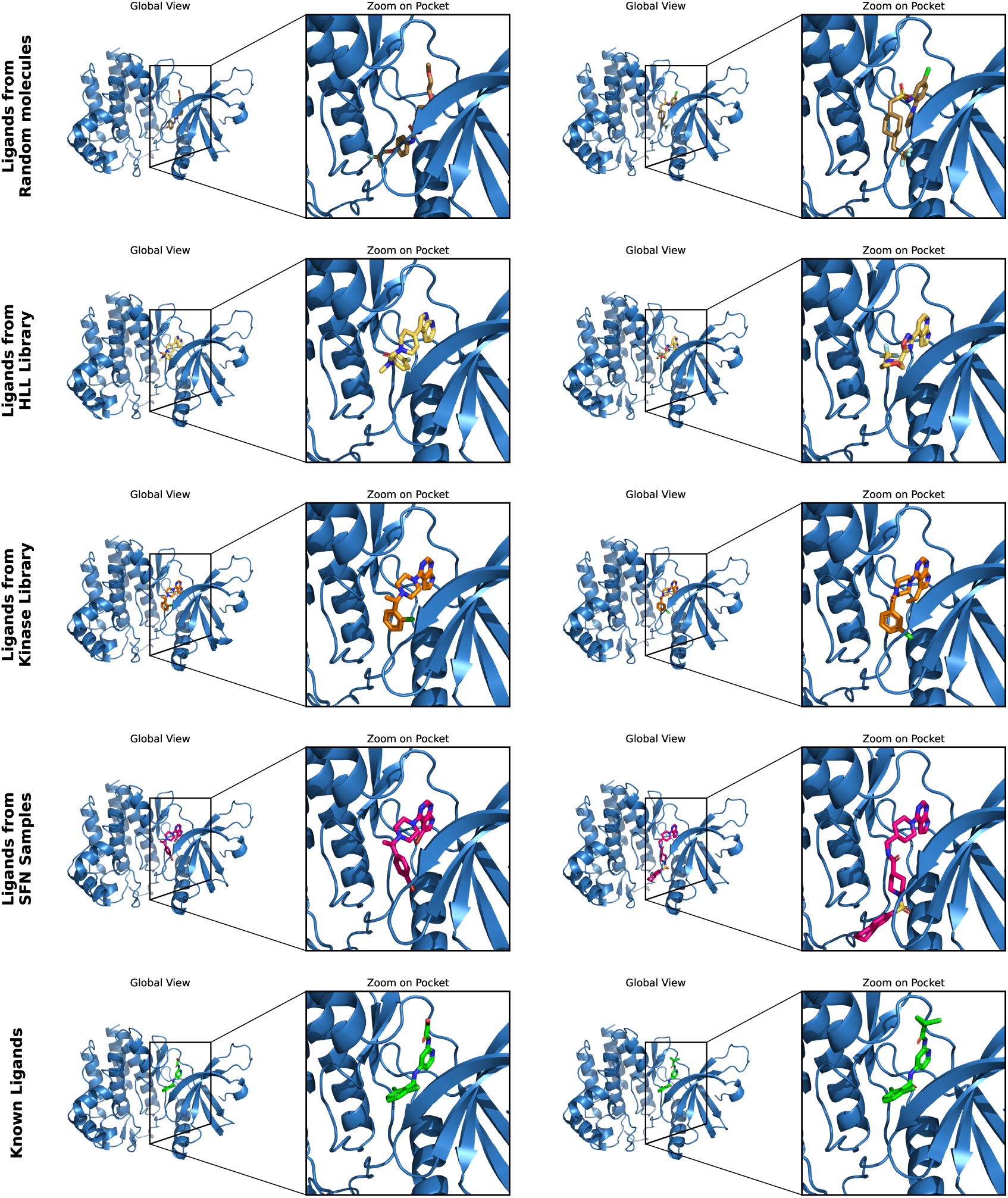
Visualization of the binding pose of the top two ligands found by each virtual screening method, as ranked by ABFE with the TYK2 protein in blue. *Note: We remind the reader that the molecules were solely optimized for their Boltz-2 score. Other properties, such as toxicity, solubility, metabolism, etc. are ignored*.

**Figure 17:**
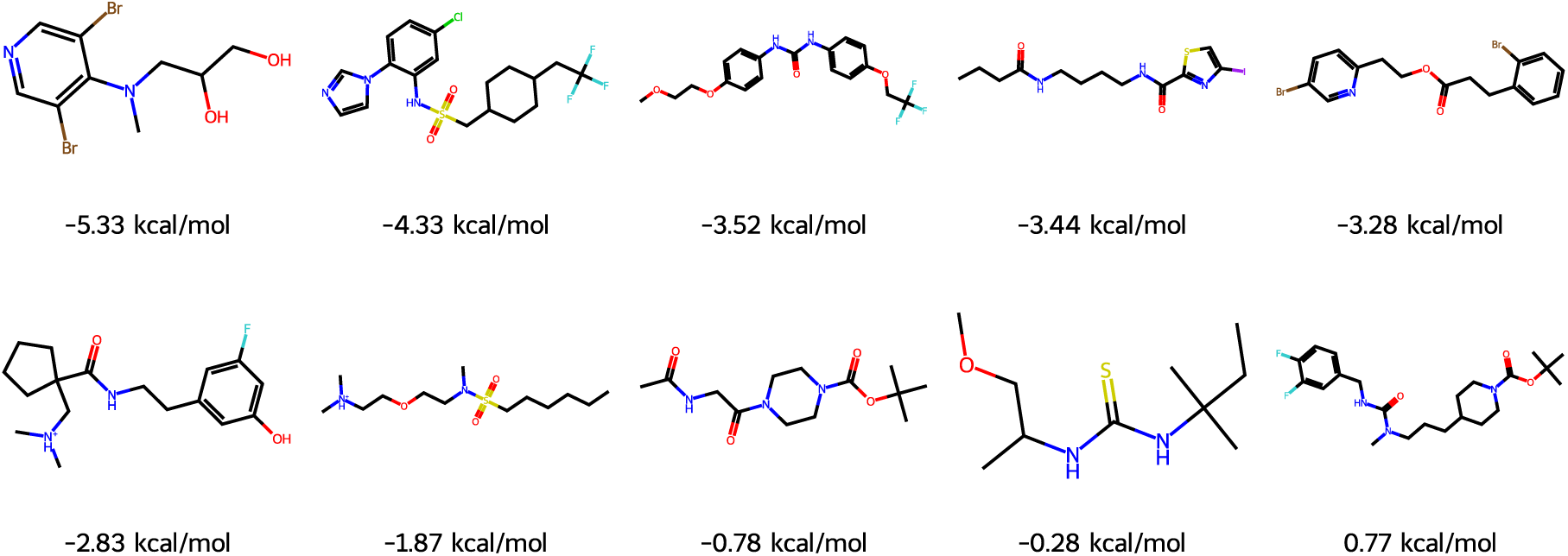
Random molecules selected for ABFE evaluation.

**Figure 18:**
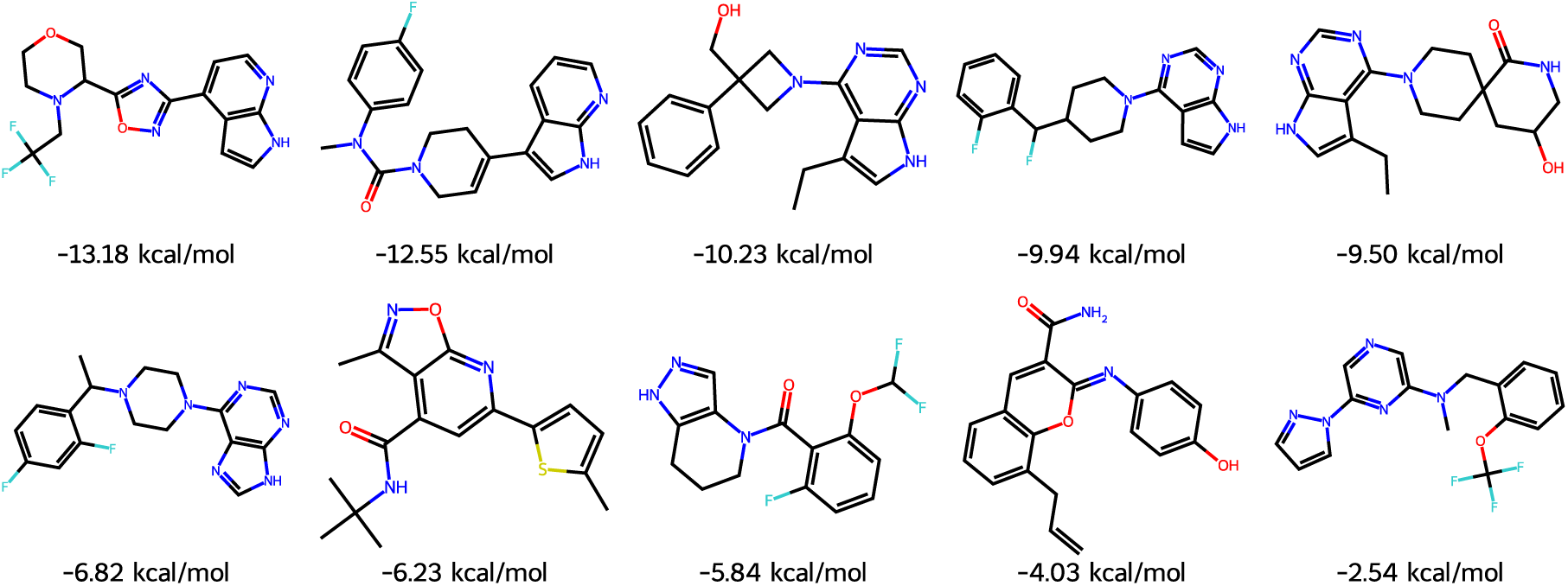
Molecules from the HLL library selected for ABFE evaluation.

**Figure 19:**
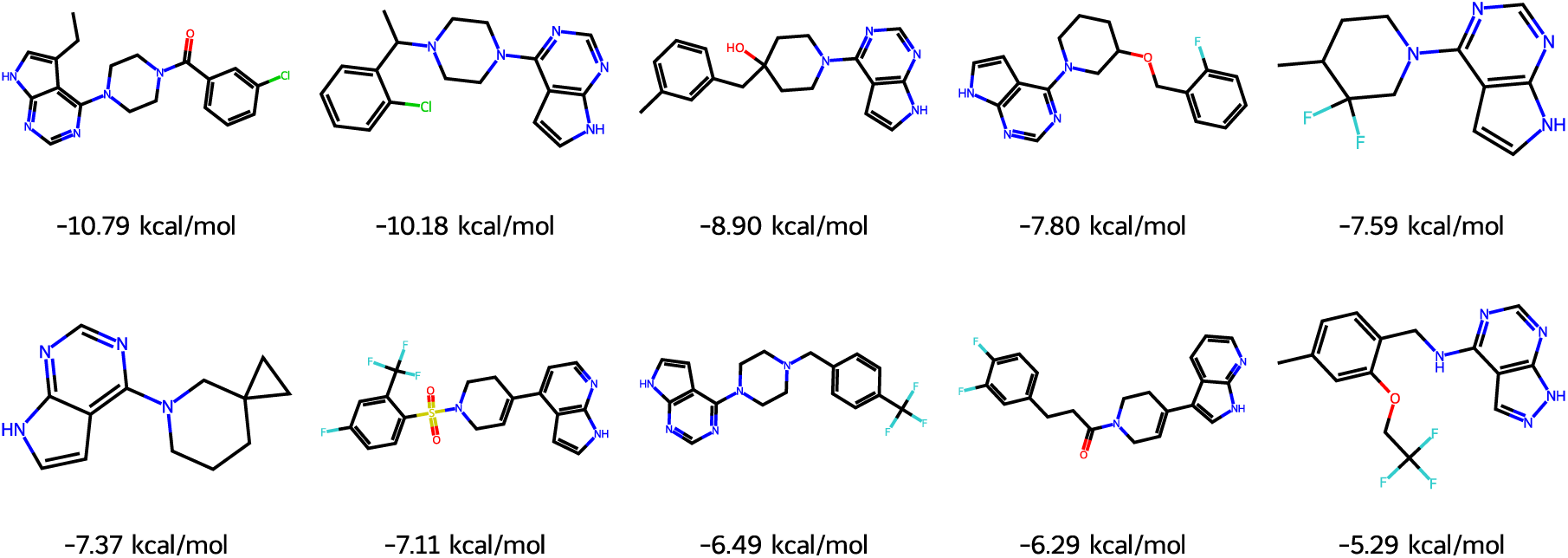
Molecules from the Kinase library selected for ABFE evaluation.

**Figure 20:**
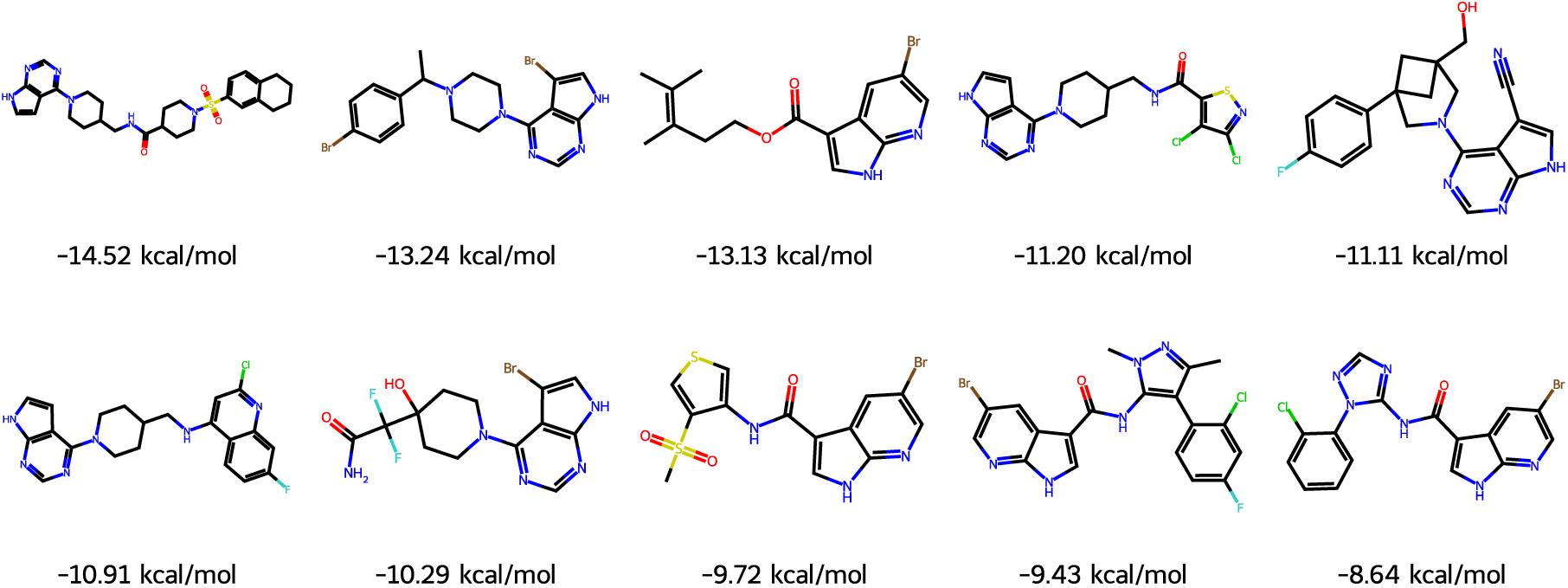
Molecules generated by SynFlowNet for ABFE evaluation. *Note: We remind the reader that the molecules were solely optimized for their Boltz-2 score. All other properties, such as toxicity, solubility, metabolism, etc. are ignored*.

##### E.3.2 ABFE validation protocol

Before ABFE evaluation, we select the most dominant tautomer state at pH=7.4 using ChemAxon. Subsequently, we use Boltz-2 to co-fold the selected compounds with TYK2 and call our Boltz-ABFE protocol to estimate ABFE values. For compounds with multiple stereoisomers, we select the isomer with the larger ABFE value.

##### E.3.3 Ligands visualisations

##### E.3.4 Similarity analysis to public TYK2-binders

The Boltz-2 structure module was trained using TYK2 protein-ligand complexes from the PDB. To assess whether Boltz-2 successfully generalized within the ligand space to steer the generation of novel candidate TYK2 binders, we collected the 47 public TYK2 inhibitors from the KLIFS database [Kanev et al., 2021] that correspond to the co-crystalized TYK2 inhibitors from the PDB. Then, we computed their Morgan fingerprint Tanimoto similarity to the ABFE-tested compounds generated by SynFlowNet. The similarity scores were computed both on the Murcko scaffolds and for the full molecules for each compounds pair (in Figure 22 we show the similarity matrix between scaffold pairs since this is the most restrictive metric). As shown in Figure 22, none of the SynFlowNet–KLIFS ligand pairs exhibited high similarity, with a maximum Tanimoto score of just 0.396 between the most similar scaffold pairs. In Figure 23, we show the most similar KLIFS ligand for every ABFE-tested ligand generated by SynFlowNet. We observe that the model captured the hinge-binding relevance of a pyrrolopyrimidine-like heterocycle, a well-established motif in orthosteric kinase inhibitor design. However, it reuses this scaffold across a range of diverse chemotypes that remain structurally distinct from their closest KLIFS counterparts.

**Figure 21:**
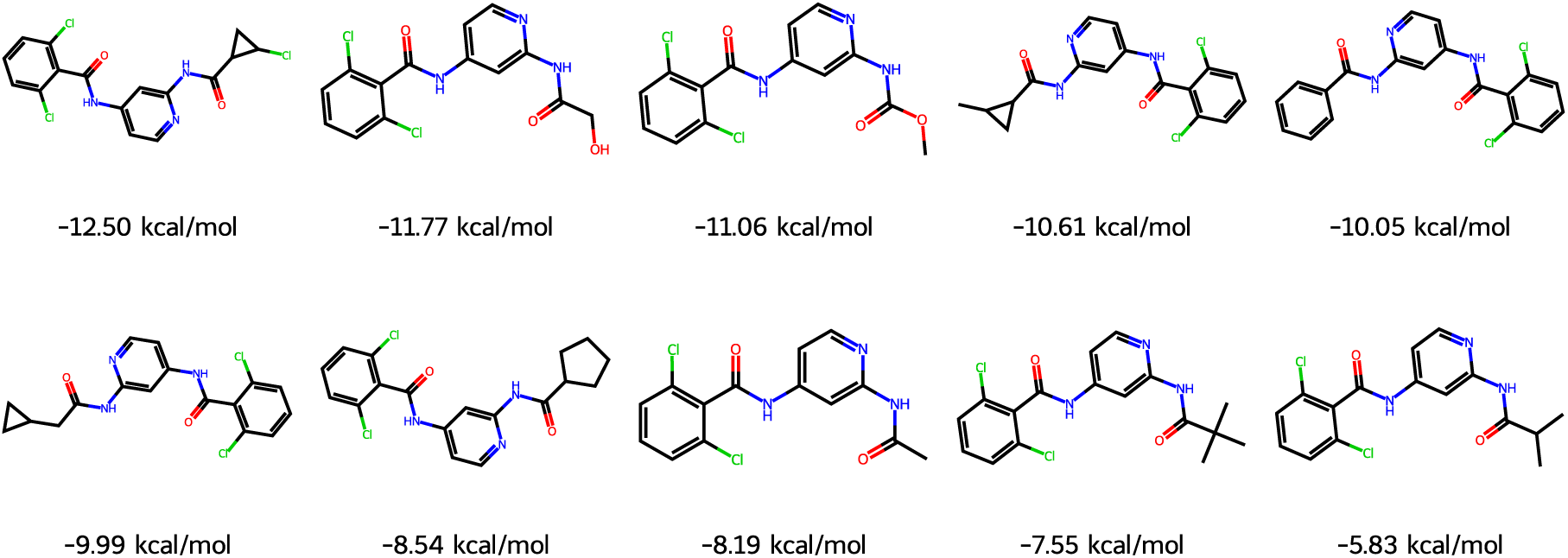
Public TYK2 binders from the protein-ligand benchmark selected for ABFE evaluation.

**Figure 22:**
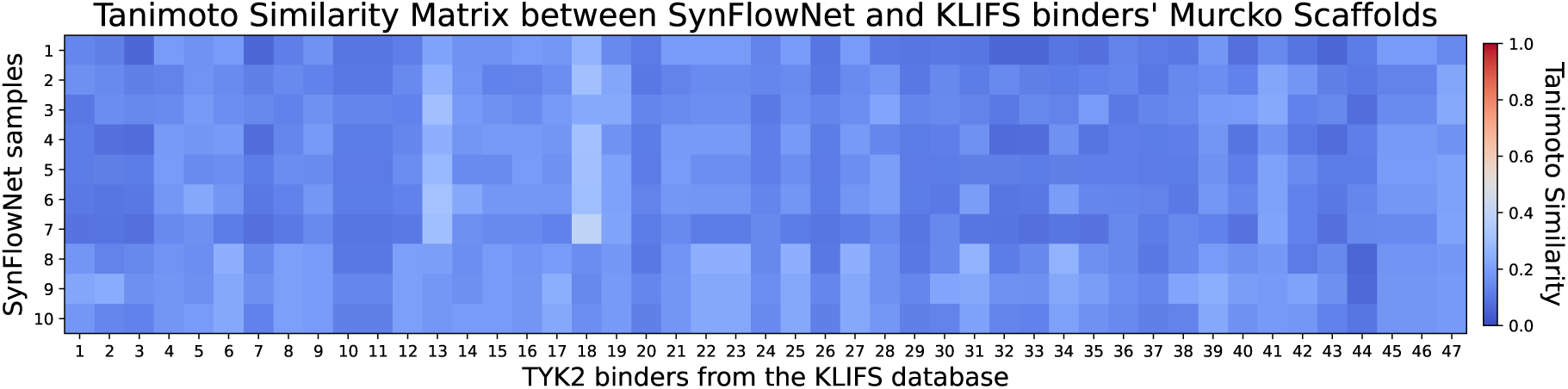
Similarity matrix between the 10 generated compounds from the SynFlowNet screen, and the TYK2 binders from the PDB as obtained from the KLIFS database. The generated compounds show significant novelty, with at most 0.396 Tanimoto similarity with known binders.

**Figure 23:**
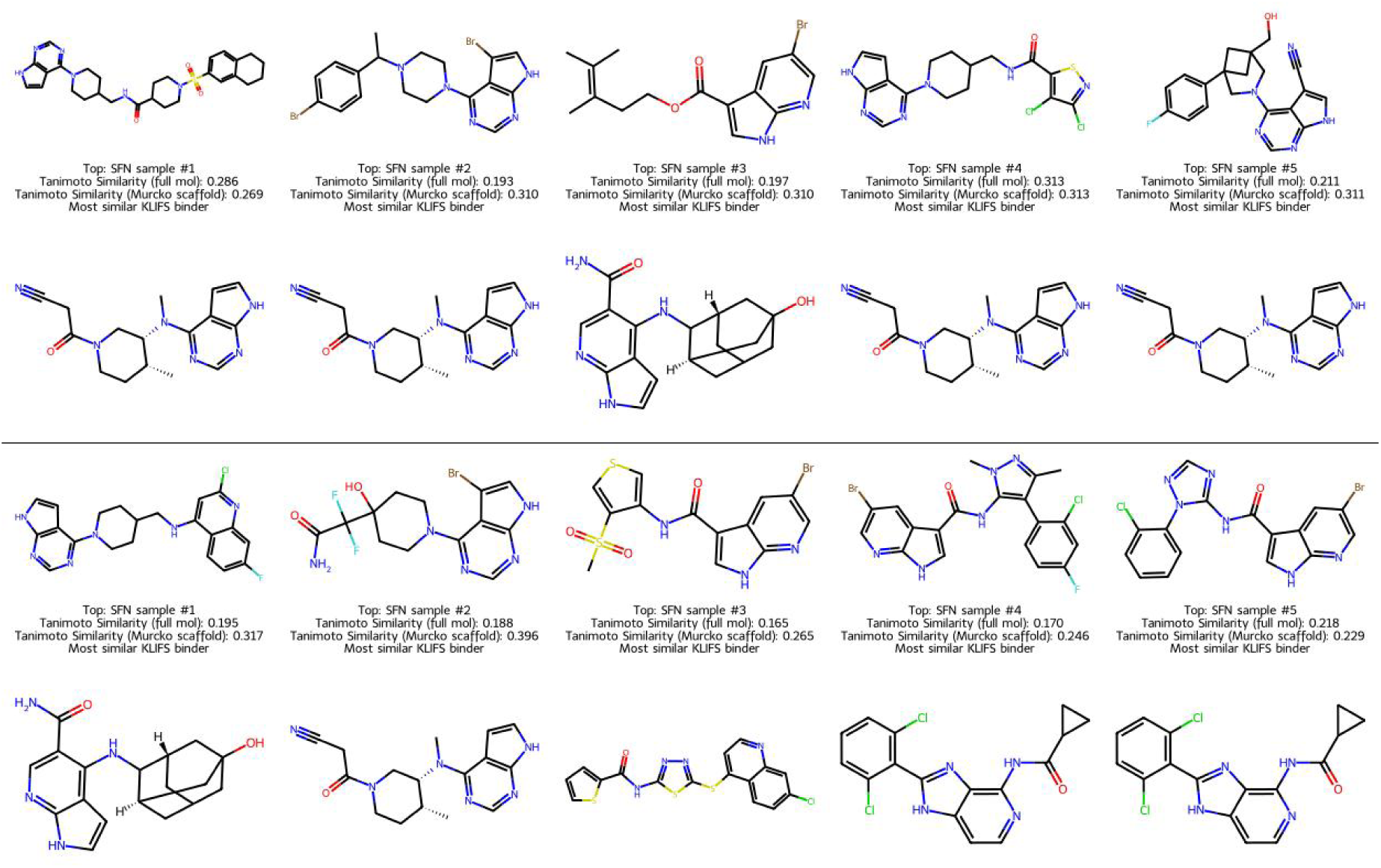
For each generated ligand from our SynFlowNet model, we show the most similar TYK2 binder from the PDB as curated in the KLIFS database. Although we find some similar structural groups, the generated ligands exhibit significant novelty throughout their entire structures, encompassing both the scaffolds and their decorations. *Note: We remind the reader that the molecules were solely optimized for their Boltz-2 score. Other properties, such as toxicity, solubility, metabolism, etc. are ignored*.

## Footnotes

1 Code, weights and data available at https://github.com/jwohlwend/boltz.

2 Preprint available soon.

